# Human single-neuron correlates of somatosensation

**DOI:** 10.64898/2026.10.07.757446

**Authors:** Bingchuan Liu, David A. Bjånes, Liang She, Kelsie Pejsa, Brian Lee, Charles Y. Liu, Richard A. Andersen

## Abstract

Understanding the neural correlates of consciousness poses a central challenge in neuroscience, with longstanding debate over which cortical areas underlie subjective sensory perception^1–3^. In the tactile domain, posterior parietal cortex (PPC) and primary somatosensory cortex (S1) are candidate cortical substrates for neural correlates of consciousness due to their crucial roles in somatosensory processing^4–8^. Yet whether and how these regions contribute to conscious tactile perception remains unclear^9–11^. Here, to address this issue, we leveraged chronic intracortical recordings to simultaneously monitor large ensembles of single neurons in grasp-related cortical areas from three participants with tetraplegia while natural vibrotactile stimuli were delivered to a sensate region of the upper limb. We demonstrate that an orchestration of neural activity emerged across the grasp-related parietal-frontal network at near-threshold perception. Within this distributed network, we then focused on PPC and S1. While S1 responses closely tracked the amplitude of the physical stimulus, PPC activity covaried more strongly with awareness and confidence. This single-neuron evidence reveals PPC and S1 have complementary neural codes with PPC transforming physical stimulation into conscious perception whereas S1 is more sensitive to stimulus amplitude.

---

Touch is omnipresent in human experience. It plays a crucial role in development and social interaction, underpinning activities of daily living. Across the human lifespan, tactile feedback guides everyday actions, from pressing a smartphone screen and typing on a keyboard to safely manipulating objects in the world. Despite its ubiquity and importance, the neural basis of the subjective perception of touch remains elusive^1,2,12^. Where in the brain does conscious tactile perception emerge?

The neural correlate of consciousness, defined as the minimal neural mechanisms collectively sufficient for a conscious percept^1,2,13^, offers a principled approach to localizing this substrate of conscious sensory input. A growing body of evidence places the neural correlate of consciousness within a temporo-parietal-occipital zone of posterior cortex in humans^1,3,14,15^. In the somatosensory domain, posterior parietal cortex (PPC) is of particular relevance because of its role in sensory and motor integration^6,7^. PPC lies within this posterior zone.

There has been ongoing debate about whether neural activity in these regions correlate with conscious somatosensory perception, especially in primary somatosensory cortex (S1), which represents early afferent input^4,5^. Some accounts contend that S1 primarily reflects early input without generating awareness, showing similar evoked responses for perceived and missed stimuli in monkeys^9^, echoing evidence from vision that sensory signals can be processed at early stages without necessarily being consciously perceived^16,17^. Conversely, transcranial magnetic stimulation of S1 can impair tactile detection^18^, while right parietal lesions can disrupt tactile awareness, producing neglect or extinction^19,20^. Recent human intracortical recordings further show that imagined somatosensory percepts activate PPC more robustly than S1^21^. Thus, whether and how human PPC and S1 jointly support tactile awareness remains an open question that requires direct comparison of neural activity in both regions during perceptual reports.

Addressing this question necessitates sufficient spatiotemporal precision to capture perceptual processing across cortical regions. However, electroencephalography (EEG) and magnetoencephalography (MEG) lack spatial specificity, whereas functional magnetic resonance imaging (fMRI) provides only an indirect measure of neural activity through hemodynamic signals, making it difficult to resolve the underlying neural processes. Single-neuron recordings offer superior spatiotemporal precision, but most work has been in nonhuman primates and cannot yield explicit subjective reports^9,22^. A few human single-neuron recording studies have begun to reveal the neural correlates of tactile consciousness^23^. These studies have identified perception-sensitive neurons in the subthalamic nucleus and thalamus^24^, sustained threshold-sensitive responses in posterior perisylvian cortex^25^ and evidence accumulation in PPC relating to both detection and confidence^11^. Extending this nascent line of research, we provide the first simultaneous recordings across S1 and PPC regions in humans to directly compare their roles in tactile perception.

We leveraged the rare opportunity provided by chronic intracortical recordings in three participants with tetraplegia enrolled in a clinical trial to examine how neurons encode tactile perception in humans. Because tactile information is essential for grasping and object manipulation, we recorded single-neuron activity simultaneously from grasp-related cortical areas, including PPC, S1, the primary motor cortex (M1) and ventral premotor cortex (PMv), as well as from the dorsolateral prefrontal cortex (dlPFC). In this study we focus on S1 and two PPC regions: the supramarginal gyrus (SMG), which contributes to somatosensory processing and sensorimotor integration^21,26^, and anterior intraparietal area (AIP), which supports somatosensory-guided grasping and sensorimotor transformation^27,28^. Participants performed near-threshold vibrotactile detection tasks where identical stimuli were sometimes felt and sometimes missed along with subjective confidence ratings, and a ramping task to assess amplitude modulation. This design guarantees a sufficient number of trials for each perceptual condition and enables separation of physical input from perceptual awareness and subjective confidence on trials with identical stimulation. We hypothesized that PPC and S1 would contribute to conscious subjective perception through distinct neural representations (Extended Data Fig. 1a).

Our large-scale simultaneous single-neuron recordings from PPC and S1 in three participants revealed a functional dissociation in how these cortical areas encode tactile sensations. First, we observed that near-threshold vibrotactile perception was accompanied by circuit-wide activity across the cortical grasp network. Second, we demonstrate that PPC preferentially encodes perceptual awareness and confidence of touch whereas S1 more robustly represents stimulus amplitude, suggesting a functional specialization across cortical stages of somatosensory processing. Together, these findings delineate complementary roles of PPC and S1 in cortical somatosensory processing, providing significant insights into the neural basis of subjective tactile perception and having important implications for basic and clinical neuroscience, as well as for sensory brain–computer interfaces aimed at restoring touch perception.

## Results

### Task, recording and distributed coding in the grasp network

To identify implantation sites, we used fMRI to localize grasp-related cortical regions^29,30^ in three participants with tetraplegia (Participants 2–4). Microelectrode arrays (4 or 6 per participant, comprising 288 or 384 channels in total) were implanted in the PPC (SMG and AIP), S1, M1, PMv, and dlPFC (Fig. 1a). Neural activity was recorded simultaneously across these regions during vibrotactile stimulation tasks (Fig. 1b). Vibrotactile stimuli were delivered using a tactor armband placed on a sensate region of the contralateral limb (Fig. 1c), which varied by participant (Participant 2: elbow, finger, or bicep; Participant 3: bicep; Participant 4: finger or bicep; Extended Data Fig. 4a,b). After each trial, the participants reported their subjective sensory perception (felt or non-felt) and confidence ratings (from 1 to 10) (Fig. 1d). A proportion of catch trials with no vibrotactile stimulation was included in each block, and trials can be classified as hits or misses when the stimulus is present and as false alarms or correct rejections when absent (Extended Data Fig. 1b). Participants first completed a one-up three-down procedure to estimate their daily individual vibrotactile detection threshold (Fig. 1e; example psychometric curve in Fig. 1f) and then conducted a near-threshold task using the estimated threshold (Extended Data Fig. 1c). In a separate ramping task, vibrotactile stimuli were placed on one of the sensate regions (elbow for Participant 2, bicep for Participant 3, finger for Participant 4) and presented in random order from a predefined set of amplitudes to sample a broad intensity range (Extended Data Figs. 1d and 7a).

**Fig. 1.**
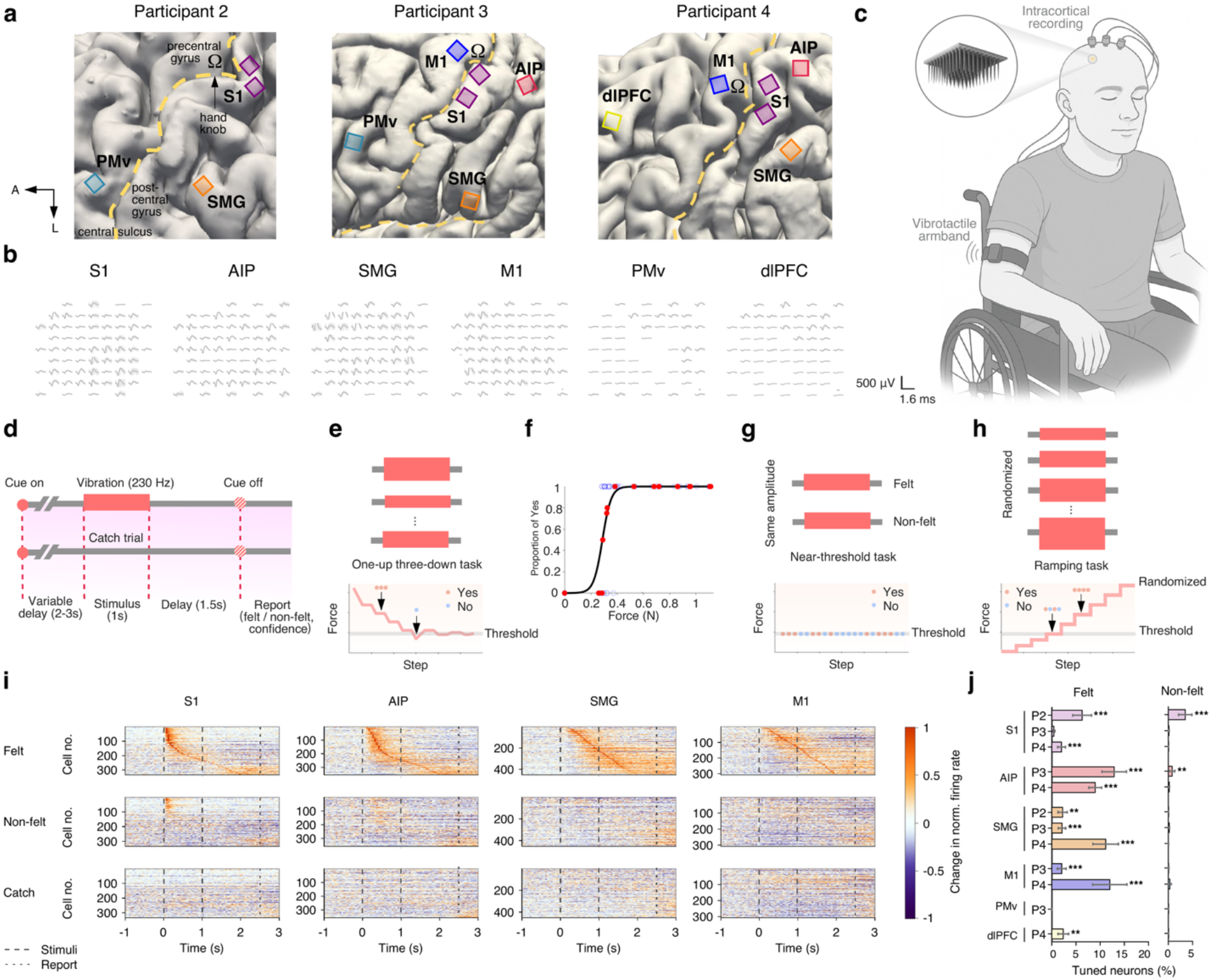
Task design, behaviour and single-neuron responses across cortical areas. **a,** Implant locations in three participants. Four arrays were implanted in Participant 2, and six arrays in each of Participants 3 and 4. Data analyses focused on arrays placed in the posterior parietal cortex (PPC) and primary somatosensory cortex (S1). **b,** Example spike-sorted neuronal activity. Representative spike waveforms from 6 different areas implanted. **c,** Experimental design. **d,** Trial structure of the vibrotactile task. **e,** One-up three-down staircase procedure. **f,** Example psychometric detection curve from the one-up three-down procedure. The x axis shows the force measured by an accelerometer attached to the tactor, and the y axis shows the proportion of yes responses. **g,** Schematic of near-threshold stimulation. After determining each participant’s detection threshold, identical vibrotactile stimuli were delivered at that level, resulting in variable perceptual outcomes where the same stimulus was sometimes felt and sometimes not. **h,** Schematic of the ramping task. **i,** Normalized firing rate changes for felt, non-felt and catch trial conditions in representative cortical areas. The x-axis denotes time from stimulus onset and the y-axis individual neurons, which were sorted by response latency and pooled across three participants. Yellow indicates increased and purple indicates decreased firing rates relative to baseline (-2 to 0 s), with dashed lines marking the stimulus period and the report onset. **j,** Proportion of tuned neurons during felt and non-felt near-threshold trials across participants (P2, Participant 2; P3, Participant 3; P4, Participant 4). Bars show the mean proportion of tuned neurons across sessions in each area, with error bars indicating the standard error of the mean (s.e.m.). Asterisks indicate that the observed proportion of tuned neurons exceeded the permutation-derived null distribution (* *P* < 0.05, ** *P* < 0.01, *** *P* < 0.001). Further details are provided in the Supplementary Information.

In the near-threshold task (Fig. 1g), a total of 61 sessions were collected across the three participants, with an average of 107.8 ± 5.2 trials per session. Across all sessions, we recorded from 17,916 single neurons spanning six cortical areas: S1 (*n* = 6,290), AIP (*n* = 2,735), SMG (*n* = 4,786), M1 (*n* = 3,031), PMv (*n* = 436), and dlPFC (*n* = 638). Since spike sorting was performed independently for each session, we assessed potential cross-session overlap of recorded neurons and found generally low putative matching rates across sessions (Participant 3: median 5.08%; Participant 4: median 6.29%; Extended Data Fig. 9h–j). The actual input force did not differ significantly between felt and non-felt trials (two-tailed rank-sum test, *P* = 0.86, Extended Data Figs. 1g and 4c,d). In the separate ramping task (Fig. 1h), 18 sessions were recorded from the same three participants, yielding 5,768 single neurons: S1 (*n* = 1,747), AIP (*n* = 965), SMG (*n* = 1,566), M1 (*n* = 1,074), PMv (*n* = 181), and dlPFC (*n* = 235).

We first asked whether subjective tactile awareness and tactile-evoked activity are represented in a single cortical area or instead recruits a distributed circuit. Area S1 was the only area that showed substantial evoked responses below threshold, and when the stimulus was felt, widespread activity was seen across the cortical areas of the grasp network in a broad set of channels (Fig. 1i,j; Extended Data Figs. 3a and 5a,c–g). Strikingly, we found S1, PPC, M1, and dlPFC exhibited significantly more neurons that carried tactile awareness information than expected from a shuffled distribution (permutation test, *P* < 0.05; Extended Data Fig. 3b), with the strongest recruitment in PPC and M1, weaker recruitment in S1 and smaller populations in prefrontal regions (Extended Data Figs. 3a and 5a). In the ramping task, supra-threshold vibrotactile stimulation significantly modulated neurons across all recorded areas (permutation test, *P* < 0.05; Extended Data Fig. 2c). The neuron counts were highest in S1 (*n* = 555), PPC (AIP, *n* = 548; SMG, *n* = 272) and M1 (*n* = 282), whereas fewer modulated neurons were recruited in PMv (*n* = 25) and dlPFC (*n* = 8) (Extended Data Fig. 2d,f). Across areas, tactile stimulation gave rise to both excitation and inhibition of single neurons’ firing rate (Extended Data Fig. 2b,e,g,h). Most regions showed predominantly excitatory responses (Extended Data Fig. 2e,g), indicating that supra-threshold touch engages strong activation throughout the cortical grasp network.

Within the distributed network, we next asked whether perception on individual trials was invariably accompanied by activation of all recorded areas or could instead occur with different subsets of the network. Single-trial analysis revealed significant activation across multiple combinations of cortical areas during felt trials, with no single combination shared by all perceptual trials (Extended Data Fig. 10a). Thus, activation of every recorded area was not obligatory for perception on individual trials. However, for some area combinations, co-activation probabilities significantly exceeded the products of individual activation probabilities (Extended Data Fig. 10b), suggesting that these area combinations were not simply explained by independent activation of individual regions (example single-trial activity in Extended Data Fig. 10c). Together, these results indicate that tactile perception is associated with coordinated recruitment of different subsets of the grasp network across individual perceptual trials.

### Enhanced neuronal correlate for subjective tactile awareness in PPC relative to S1

Within the distributed cortical network recruited by near-threshold touch, we next focused on PPC and S1 to characterize the neuronal correlates of subjective tactile awareness. In both regions, responses averaged across neurons diverged between felt and non-felt trials, and this effect was also evident at the level of individual neurons (Fig. 2a,b). During catch trials without stimulation, these neurons also showed prominent activation on false alarm trials, with activity patterns resembling those elicited by actual vibration (Fig. 2b,c), indicating that cortical representations of sensory awareness can emerge even in the absence of external input. To control for the potential confound of explicit perceptual reporting, we further performed a no-report control task^31^ using the same procedure as the near-threshold task, except that participants were not required to provide a behavioural report (Extended Data Fig. 9f). Representative PPC and S1 neurons showed similar responses in report and no-report blocks, suggesting that these neural responses persisted in the absence of overt reporting and were unlikely to be explained solely by report-related motor activity (Extended Data Fig. 9g). Notably, as shown in Figs. 1i and 2a, averaged firing rates showed that S1 was active for stimuli that were below the perceptual threshold, whereas the other brain areas showed substantial activation only above perceptual threshold. This result implies that S1 is not the site of sensory awareness, and further suggests that awareness is a property of the simultaneous activation of a group of grasp network areas downstream of S1.

**Fig. 2.**
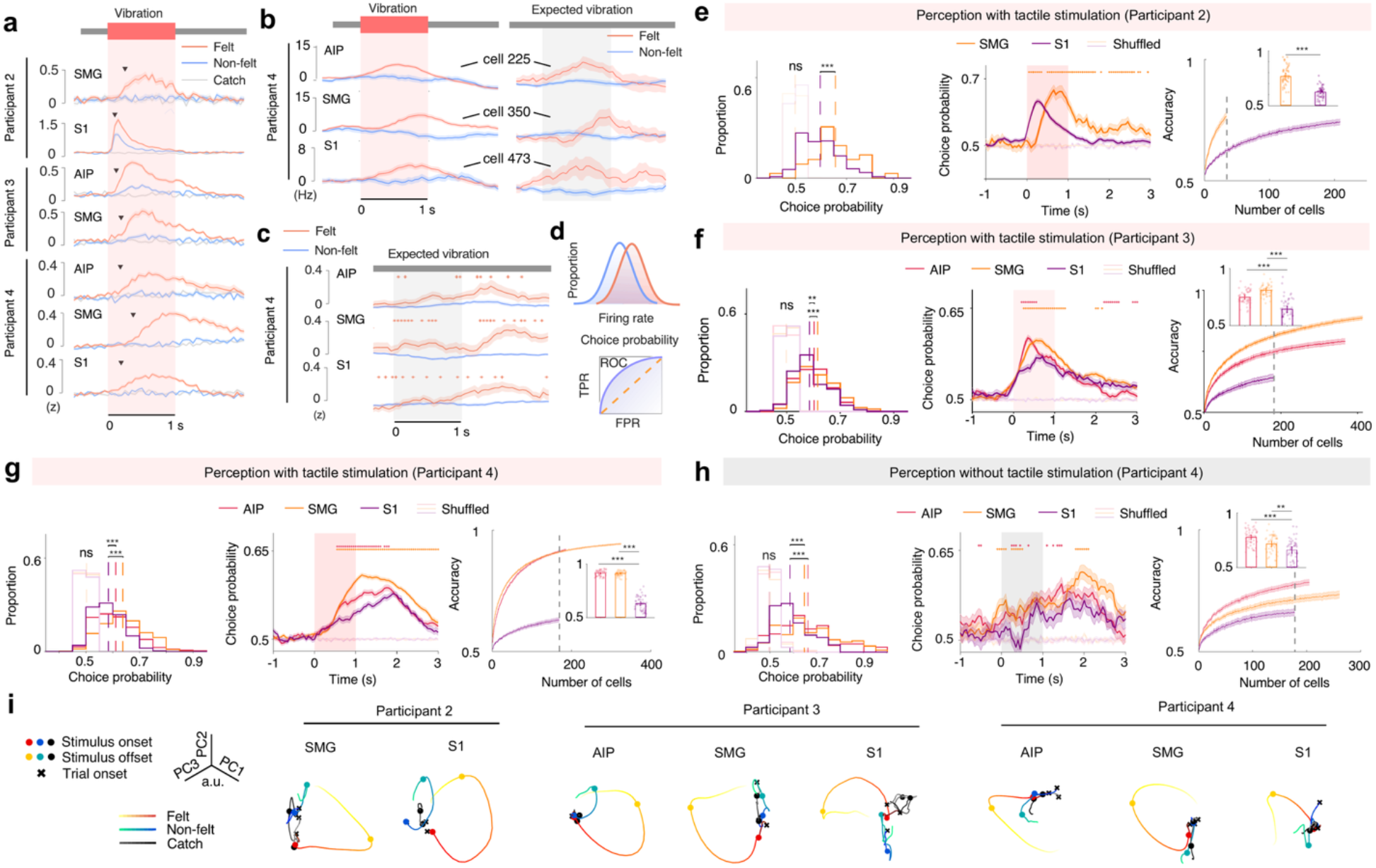
PPC and S1 are differentially modulated by subjective tactile awareness. **a,** Average firing dynamics for felt, non-felt, and catch trials in PPC and S1. Arrows mark the first time bins where felt responses differed significantly from non-felt responses. **b,** Example neuronal responses in Participant 4 during actual and catch trials. **c,** Average normalized population firing rates during catch trials in Participant 4. Asterisks indicate significant differences. **d,** Illustration of choice probability. TPR: true positive rate; FPR: false positive rate. **e**–**g,** Choice probability and population decoding during vibration. Left panels: Distributions of choice probability. Middle panels: Choice probability over time. Right panels: Neuron-dropping analysis. Inset shows accuracy at the neuron count marked by the dashed line. PPC showed higher choice probability than S1 (Participant 2: *P*_SMG,S1_ = 8 × 10^−5^; Participant 3: *P*_SMG,S1_ = 1.1 × 10^−5^, *P*_AIP,S1_ = 0.0038; Participant 4: *P*_SMG,S1_ = 3.0 × 10^−18^, *P*_AIP,S1_ = 3.3 × 10^−4^; two-tailed rank-sum test), and outperformed S1 at matched neuron count (Participant 2: *P*_SMG,S1_ = 3.2 × 10^−12^; Participant 3: *P*_SMG,S1_ = 1.5 × 10^−13^, *P*_AIP,S1_ = 1.9 × 10^−8^; Participant 4: *P*_SMG,S1_ = 7.1 × 10^−18^, *P*_AIP,S1_ = 7.1 × 10^−18^; two-tailed rank-sum test). Asterisks indicate significant differences between PPC and S1. Neuronal sample sizes are in the Supplementary Information. **h,** Choice probability and population decoding during catch trials in Participant 4. PPC showed higher choice probability than S1 for distinguishing catch felt from catch non-felt trials (*P*_SMG,S1_ = 1.6 × 10^−5^, *P*_AIP,S1_ = 2.4 × 10^−4^; two-tailed rank-sum test) and higher LDA decoding accuracy (*P*_SMG,S1_ = 2.8 × 10^−3^, *P*_AIP,S1_ = 8 × 10^−8^; two-tailed rank-sum test). **i,** Neural dynamics for felt, non-felt, and catch conditions across participants. Warmer colours represent felt trials, cooler colours represent non-felt trials. Black represents catch trials.

We next compared PPC and S1 in their sensitivity to subjective sensory awareness. To quantify how well individual neuronal activity predicted perceptual choice, we computed the choice probability for each neuron. This metric reflects the separability of firing rate distributions between felt and non-felt trials (Fig. 2d). Across the three participants, SMG or AIP neurons consistently showed significantly higher choice probabilities than S1 during the analysis window (Fig. 2e–g; multi-region extension in Extended Data Fig. 3c,d), indicating stronger perceptual encoding within PPC, although individual differences were present in the onset and temporal profile of choice probability. In participants with matched receptive fields, felt–non-felt divergence showed a trend toward earlier onset in S1 than in the recorded PPC regions (Extended Data Fig. 8a). Thus, earlier divergence in S1 coexisted with stronger perceptual code in PPC. This PPC advantage persisted after excluding putatively overlapping neurons across sessions (Extended Data Fig. 3j,k). For multi-area combinations, choice probability was generally higher than for individual areas, further supporting greater perceptual information in distributed activity (Extended Data Fig. 10d). The same trend appeared in near-threshold trials of the ramping task (Extended Data Fig. 7d). Across stimulation locations, the proportion of perception-sensitive activations varied more strongly in S1 than in PPC, and choice probability differed significantly across locations and participants in S1 but not in PPC (Extended Data Fig. 4e,f). Elevated choice probabilities in PPC relative to S1 remained consistent during false alarm trials despite the absence of physical stimulation (Fig. 2h).

To further assess population-level encoding, we performed a neuron-dropping analysis using a linear discriminant analysis (LDA) classifier trained to distinguish felt versus non-felt trials. We constructed neural pseudo-populations^32^ by pooling neurons with near-threshold perceptual sensitivity across sessions within each participant. For a matched number of neurons, decoding accuracy was consistently higher in AIP or SMG compared with S1 across participants (Fig. 2e–g; multi-region extension in Extended Data Fig. 3e,f).

This PPC advantage was also observed across all tested stimulation locations, including three sites in Participant 2 and two sites in Participant 4 (Extended Data Fig. 4g,h). This advantage also extended to false alarm trials (Fig. 2h and Extended Data Fig. 3g). Moreover, cross-condition decoding showed that classifiers trained on stimulus-present hit versus miss trials generalized to stimulus-absent false alarm versus correct rejection trials, with stronger generalization in PPC and other non-S1 areas than in S1 (Extended Data Fig. 3h,i). Consistent with this perceptual advantage, behavioural choice in the vibrotactile discrimination task was decoded more accurately from AIP and SMG than from S1 when physical force and neuron count were matched (Extended Data Fig. 9e). These population-decoding findings further corroborate that PPC exhibits greater perceptual sensitivity than S1 during near-threshold stimulation.

To examine the neural dynamics underlying subjective perception, we projected neural activity from pseudo-populations in PPC and S1 into a three-dimensional state space using principal component analysis (PCA). In all three participants, trajectories corresponding to felt and non-felt trials diverged clearly, indicating that subjective sensory awareness is reflected in distinct population-level dynamics within a low-dimensional neural state space (Fig. 2i). To further compare population-level geometry across cortical areas, we aligned low-dimensional neural trajectories using Procrustes analysis. M1 and S1 showed more similar neural manifold structure, whereas AIP and SMG were more closely aligned with each other, suggesting a distinction between sensorimotor and higher-order parietal representations (Extended Data Fig. 8c–f). To quantify the temporal stability of these perceptual dynamics, cross-time representational similarity analysis (RSA) further showed that perceptual structure was more sustained in PPC than in S1, suggesting that higher-order parietal areas contribute to the maintenance of tactile perceptual representations beyond early sensory encoding (Extended Data Fig. 8g–i).

### Modulation by physical stimulus amplitude is stronger in S1 than in PPC

Having used the near-threshold task to isolate conscious tactile perception under matched physical input, we next used the ramping task^25^ to characterize physical intensity coding across the same cortical network. Vibrotactile stimuli of nine amplitudes were randomly presented, including a catch trial with no stimulation (-100%), two sub-threshold levels (Sub1, -75%; Sub2, -50%), one threshold level (0%), and five supra-threshold levels (Supra1–Supra5: +20%, +40%, +60%, +80% and +100%; Fig. 3a). Higher-amplitude inputs elicited stronger activations across the grasp network, with progressively increasing firing observed in cortical regions (Fig. 3a; M1 in Extended Data Fig. 7c). Neurons in PPC and S1 exhibited significant trial-by-trial correlations between firing rate and input force (Fig. 3b). For subsequent analyses, we focused on task-modulated neurons identified via one-way ANOVA, based on significant firing rate differences across the nine stimulus conditions. At the single-neuron level, both normalized firing rates and the area under the receiver operating characteristic curve (AUC) increased with amplitude (Fig. 3c,d), where AUC was computed by comparing the firing rate of each amplitude to that of the catch trial. Participant 3 was not used in the following amplitude-sensitivity analysis due to a lack of tactile-evoked neurons (Extended Data Fig. 2f,g), as the receptive field of the array did not overlap with somatotopic regions with intact sensation. Data from this participant were retained in the other analyses.

**Fig. 3.**
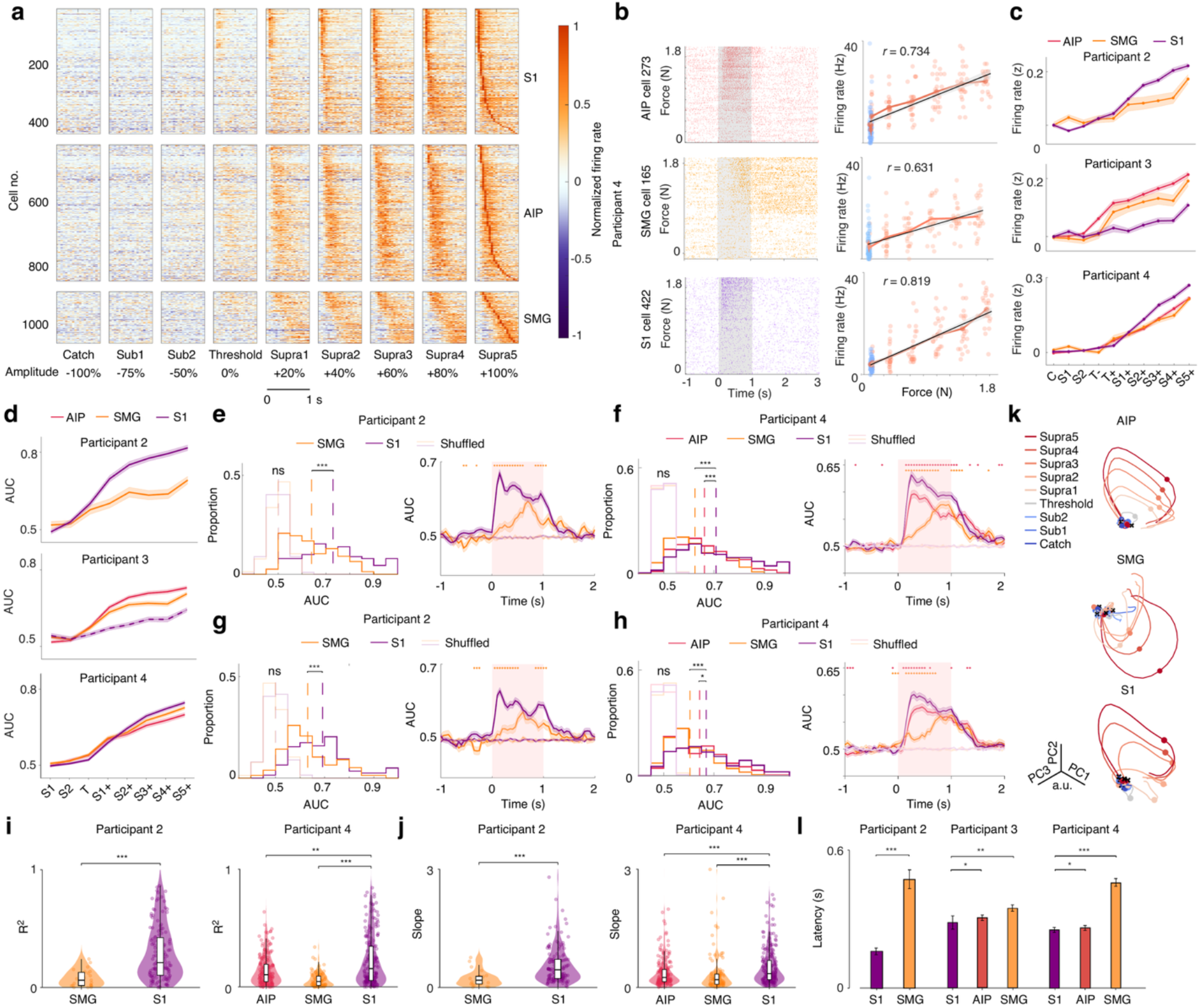
PPC and S1 encode distinct representations of physical stimulus amplitude. **a,** Neural responses to varying vibrotactile amplitudes. Normalized firing rate changes during stimulation are shown across participants for nine vibrotactile amplitudes, including five supra-threshold levels (+20%, +40%, +60%, +80%, and +100%), one threshold (0%), two sub-threshold levels (-75%, -50%), and a catch condition (-100%). **b,** Example cells from S1, AIP, and SMG showing force-dependent firing modulation. Raster plot (left) and scatter plot (right) for a representative neuron, with trials ordered by input force on the y-axis for the raster plots. In the scatter plot, red indicates felt trials and blue indicates non-felt trials. **c,** Amplitude-dependent changes in normalized firing across participants. **d,** AUC as a function of input amplitude. Data from Participant 3 were excluded from subsequent AUC analyses due to implant placement outside the sensate receptive fields, and Participant 3’s S1 is marked with a dashed line. **e**,**f,** AUC between +100% and +20% conditions. AUC was computed between +100% and +20% amplitudes for Participant 2 and Participant 4. Left panels show the distribution of AUC values for PPC and S1 neurons, and right panels show the temporal profile computed in sliding bins. **g**,**h,** AUC between +100% and +40% conditions. **i,** Coefficients of determination (R^2^) comparison between PPC and S1. S1 showed significantly higher R^2^ values than PPC (Participant 2: *P*_SMG,S1_ = 5.2 × 10^−7^; Participant 4: *P*_SMG,S1_ = 8.3 × 10^−15^, *P*_AIP,S1_ = 0.0012; two-tailed rank-sum test). **j,** Slope comparison between PPC and S1. S1 exhibited significantly steeper slopes than PPC (Participant 2: *P*_SMG,S1_ = 2.7 × 10^−5^; Participant 4: *P*_SMG,S1_ = 1.0 × 10^−7^, *P*_AIP,S1_ = 2.7 × 10^−4^; two-tailed rank-sum test). **k,** Neural dynamics across amplitude conditions in Participant 4. **l,** Neural latency for PPC and S1 during supra-threshold stimulation.

**Fig. 4.**
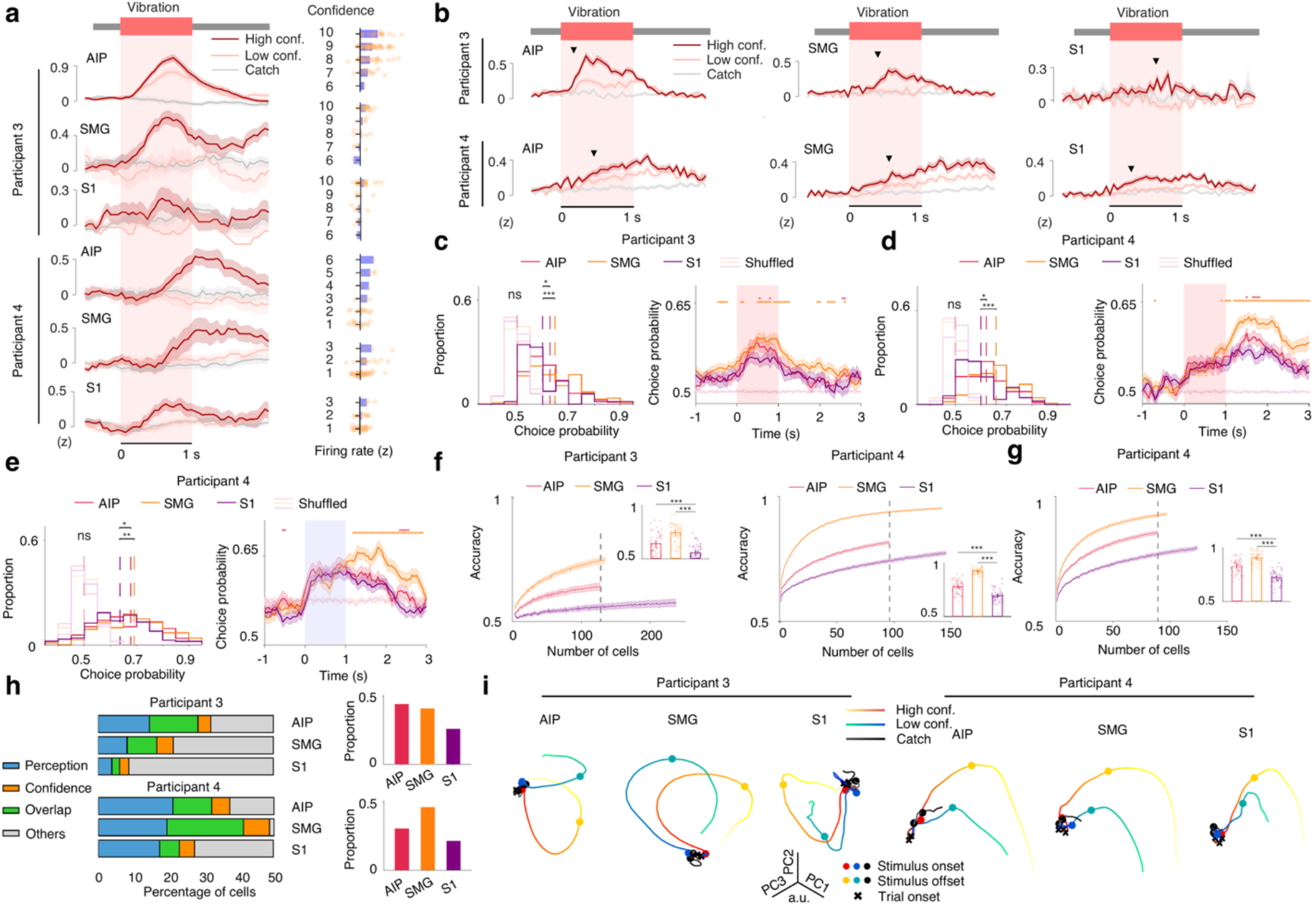
Encoding of subjective confidence in PPC and S1. **a,** Confidence dependent firing in example PPC and S1 neurons. Left panel shows normalized firing rate traces. Right panels show single-trial confidence ratings and corresponding firing rate changes for the same neurons. The y-axis indicates confidence rating and the x-axis indicates firing rate; blue bars denote trial-averaged firing rates, and dots represent individual trials. **b,** Average firing rate for high-confidence, low-confidence, and catch trials in PPC and S1. Arrows indicate time bins where high- and low-confidence trials differ significantly. **c**,**d,** Choice probability for distinguishing high-vs low-confidence felt trials. PPC showed higher choice probability than S1 (Participant 3: *P*_SMG,S1_ = 8.5 × 10^−4^, *P*_AIP,S1_ = 0.0219; Participant 4: *P*_SMG,S1_ = 9.1 × 10^−7^, *P*_AIP,S1_ = 0.0439; two-tailed rank-sum test). **e,** Choice probability for distinguishing high- vs low-confidence non-felt trials in Participant 4. The result is consistent with the felt condition, showing stronger confidence encoding in PPC (*P*_SMG,S1_ = 0.0011, *P*_AIP,S1_ = 0.0122; two-tailed rank-sum test). **f,** Neuron-dropping analysis for classifying high- vs low-confidence felt trials. PPC showed higher classification accuracy than S1 (Participant 3: *P*_SMG,S1_ = 3.5 × 10^−14^, *P*_AIP,S1_ = 3.1 × 10^−5^; Participant 4: *P*_SMG,S1_ = 1.4 × 10^−17^, *P*_AIP,S1_ = 1.4 × 10^−9^; two-tailed rank-sum test using the same neuron counts). **g,** Neuron-dropping analysis for classifying high- vs low-confidence non-felt trials in Participant 4. PPC achieved higher classification accuracy than S1 (Participant 4: *P*_SMG,S1_ = 2.3 × 10^−17^, *P*_AIP,S1_ = 1.6 × 10^−10^; two-tailed rank-sum test using the same neuron counts). **h,** Overlap between perception-tuned and confidence-tuned neurons in PPC and S1. Left panel shows the proportion of perception-tuned (blue), confidence-tuned (yellow), and overlap (mixed-selective, green) neurons among all spike-sorted units. Right panel shows the proportion of overlap neurons relative to all tuned neurons. **i,** Neural dynamics for high- vs low-confidence felt conditions.

We compared PPC and S1 in their sensitivity to changes in supra-threshold stimulus amplitude. When contrasting the highest amplitude (+100%) with lower supra-threshold levels (e.g., +20% or +40%), S1 neurons exhibited higher AUC values than PPC neurons, indicating greater sensitivity to vibrotactile amplitude (Fig. 3e–h). Although the difference in AUC decreased as the amplitude contrast became smaller, the overall trend remained consistent (Fig. 3g,h). Regression analyses between firing rate and input force showed that S1 neurons exhibited significantly higher coefficients of determination (R²) (Fig. 3i) and steeper slopes (Fig. 3j). This S1 advantage in amplitude encoding extended to non-felt trials (subliminal; Extended Data Fig. 7e), echoing stimulus-evoked S1 responses in the absence of conscious perception (Figs. 1i and 2a). To exclude the confound that subjective intensity covaries with physical amplitude (Extended Data Fig. 7b), we examined trials with matched felt-intensity ratings but different physical amplitudes, finding consistently stronger amplitude encoding in S1 than in PPC (Extended Data Fig. 7f). To control for the covariation between physical amplitude and perceived intensity, we further performed a vibrotactile two-alternative forced-choice (2AFC) amplitude discrimination task^33^ (Extended Data Fig. 1f). The analysis confirmed stronger physical force coding in S1 when behavioural choice and neuron count were matched across areas (Extended Data Fig. 9a–d). Together, these findings indicate that while both regions encode vibrotactile input strength, S1 shows a stronger modulation by amplitude variations.

To further characterize the temporal dynamics of sensory encoding, we examined the neural trajectories derived from neural pseudo-populations. Neural dynamics across amplitude levels revealed graded divergence in state space, with higher vibrotactile inputs eliciting increasingly distinct population trajectories over time in both PPC and S1 (Fig. 3k). Furthermore, we assessed neural response latency during the highest-amplitude condition (+100%), defined as the earliest time bin where firing rates significantly diverged from catch trials. As expected, S1 consistently showed shorter mean response latencies than PPC regions across participants (Participant 2: *l_S_*_1_ = 162 ms, *l_SMG_* = 491 ms; Participant 3: *l_S_*_1_ = 286 ms, *l_AIP_* = 307 ms, *l_SMG_* = 350 ms; Participant 4: *l_S_*_1_ = 254 ms, *l_AIP_* = 262 ms, *l_SMG_* = 461 ms), confirming a faster recruitment of S1 in processing strong vibrotactile inputs (Fig. 3l and Extended Data Fig. 8b).

We further combined near-threshold and ramping tasks within individual recording sessions (Extended Data Fig. 1e) to test whether perceptual-state and amplitude coding coexist within individual neurons. Neurons with near-threshold perceptual sensitivity showed stronger firing-rate correlations with amplitude and a greater likelihood of amplitude modulation, with partially overlapping perceptual-state and amplitude coding in both PPC and S1 (Extended Data Fig. 6a–c). Among neurons modulated in either task, a generalized linear model (GLM)-based modulation index indicated a stronger relative preference for amplitude coding in S1 than in SMG (Extended Data Fig. 6d–f). We identified two typical response profiles, particularly in S1 and AIP: graded ramping responses across amplitudes and nonlinear sigmoidal responses with saturation (Extended Data Fig. 6d–h). Together, these findings are consistent with a continuum of response properties that could allow cortical populations to flexibly represent perceptual state while preserving sensory information.

### Stronger neural representation of confidence in PPC than in S1

We next asked whether single-neuron activity encodes subjective confidence during near-threshold perception. The same input vibrotactile force was delivered on high- and low-confidence trials (Extended Data Fig. 1h), isolating confidence from stimulus strength. We found that many neurons from Participants 3 and 4 responded more strongly to high-than low-confidence felt trials (Fig. 4a). A significant number of neurons showed confidence tuning across levels, and this effect was reflected at the population level by significant confidence-related modulations in both PPC and S1 (Fig. 4b).

To evaluate confidence-related encoding between PPC and S1 in the near-threshold task, we assessed neural sensitivity in distinguishing high- and low-confidence felt trials. Choice probability analysis on confidence-tuned neurons revealed that neurons in PPC exhibited significantly higher choice probabilities than those in S1 (Fig. 4c,d), indicating greater single-neuron encoding of subjective confidence. These results were further supported at the population level through neuron-dropping analysis, in which classification performance for confidence levels improved more rapidly with increasing PPC neuron counts and reached higher accuracy compared with S1 (Fig. 4f). Confidence-related signals persisted even in non-felt trials, with PPC neurons showing significantly higher choice probabilities (Fig. 4e) and classification accuracy than S1 (Fig. 4g), revealing stronger confidence representation in PPC than S1 that is independent of conscious tactile detection. Consistent with this subjective coding advantage, decoding high-versus low-perceived intensity at fixed physical amplitude in the ramping task also yielded stronger perceived-intensity signals in PPC than S1 (Extended Data Fig. 7g).

Since both sensory awareness and confidence are inherently subjective, do PPC and S1 jointly encode these signatures at the single-neuron level? To address this, we investigated the neurons tuned to both confidence and near-threshold perception. Across Participants 3 and 4, a greater proportion of neurons in PPC exhibited overlapping tuning for both confidence and near-threshold perceptual awareness compared with S1 (Participant 3: SMG, 39.5%; AIP, 42.9%; S1, 25%; Participant 4: SMG, 44.7%; AIP, 29.4%; S1, 20.3%; Fig. 4h). This suggests that PPC contains neurons jointly modulated by subjective sensory awareness and the perceived certainty of sensory perception, reflecting an integrated neural code for both the detection of sensory input and the confidence associated with that perception. Furthermore, population trajectories in neural state space revealed distinct dynamics across confidence levels during near-threshold perception (Fig. 4i).

## Discussion

How subjective tactile perception emerges from neural activity remains elusive at the level of single neurons in the human brain. By delivering vibrotactile stimulation while participants performed near-threshold detection and amplitude-ramping tasks, we found that touch force is first measured early within S1, if it meets a threshold, then conscious perception occurs coincident with the engagement of a broader, distributed network that comprises cortical areas in the posterior parietal cortex and frontal lobe. These areas encompass the grasp network, which requires somatosensation for the manipulation of objects. Differences in activation exist between PPC and S1, with PPC more closely related to perception and confidence whereas S1 is more strongly related to stimulus amplitude. These findings provide further support for the hypothesis that S1 measures the tactile force, whereas PPC is part of a network responsible for perceptual awareness.

Whether the primary somatosensory cortex contributes to conscious perception has long been debated. Previous electrophysiological, neuroimaging, and stimulation studies have demonstrated that S1 activation covaries with subjective reports of tactile perception across various experimental contexts^10,34,35^. Indirect evidence from pain studies^36^ further supports the activation of S1 to the subjective perception of sensory and nociceptive events in humans. Note however, that this correlative evidence would be expected if S1 is feeding information into higher cortical areas that may be necessary for awareness. In our recordings, S1 activity also differed between felt and non-felt trials at matched stimulus force, contrasting with previous single-unit findings in monkey S1^9^. By analogy with work in vision, this modulation could reflect state-dependent variability in sensory processing^37^, including sensory gating, as well as re-entrant feedback from higher-order areas^38^.

Accumulating evidence suggests that the neural correlates of consciousness primarily reside within a temporo-parietal-occipital cortical hot zone that includes PPC and sensory areas, especially the higher-order sensory areas^1,3^. In the tactile domain, our single-neuron evidence lends support to this view by demonstrating that PPC is more closely related to perception than S1. Beyond near-threshold detection, this dissociation generalized to a discrimination task that separated physical force from behavioural choice (Extended Data Fig. 9d,e). This dissociation resonates with a fundamental hierarchical mechanism of sensory processing in somatosensation^39^ and vision^40,41^, where lower-level cortical areas tend to encode stimulus properties, whereas higher-order areas increasingly reflect perceptual choice and selectively gate behaviourally relevant sensory signals^42^. Consistent with this view, response latencies in both the near-threshold and ramping tasks showed a trend toward a temporal hierarchy from S1 through AIP to SMG, suggesting a spread of neural activity from early somatosensory cortex toward higher-order parietal areas (Extended Data Fig. 8a,b). The higher-order role of PPC is attributable to two putative somatosensory pathways^43–45^, one dedicated to tactile and object processing and the other supporting the somatosensory guidance of actions such as reaching and grasping, both receiving tactile input from S1 and together situating PPC as a hub for multisensory integration. Furthermore, consistent with previous reports of higher-order tactile somatotopy in PPC^46–48^, perceptual encoding was less location dependent in PPC than in S1 (Extended Data Fig. 4e,f), suggesting that PPC represents tactile percepts in a more abstract and less locally topographic format.

In this study, we found a shared neural correlate of touch perception across all three participants when comparing PPC and S1, despite individual differences in residual sensation, array receptive fields, and vibrotactile stimulation locations. These individual differences, together with clinically determined variation in cortical coverage, may contribute to variability in neuronal responses. Across these varied conditions, the study remained limited by its small number of participants. The limited spatial coverage of Utah arrays further restricts characterization of representations throughout each cortical area and across the somatosensory network. Despite these limitations, the consistency of our primary findings across three participants, together with extensive sampling across these different sensory and recording conditions over 100 sessions (Supplementary Table 1) and multiple control experiments addressing potential confounds, supports their robustness (Supplementary Tables 2 and 3) across the sensory conditions examined.

Beyond PPC and S1, neural responses during both supra-threshold stimulation and near-threshold perception extended across the grasp network and were especially prominent in M1, suggesting that conscious touch engages a sensorimotor circuit rather than a single cortical locus. Such network-wide encoding may support the transformation of sensory evidence into decisions and actions for interacting with the environment, through recurrent bottom-up and top-down circuits that help stabilize the fidelity of tactile feedback despite local variability in neural activity. This distributed profile parallels findings in vision from recent adversarial tests of global neuronal workspace and integrated information theories^3^, which identified conscious content in sensory as well as frontal cortex alongside sustained posterior responses, and extends these findings to touch at the level of single neurons. By revealing the single-neuron substrate of tactile perceptual signals, our findings provide critical evidence for linking single-neuron codes to systems-level theories of the neural correlates of consciousness and for guiding the design of closed-loop somatosensory brain–computer interfaces.

## Acknowledgements

We are grateful to participants FG, AN and GB for their dedication, time and invaluable contribution to this study. This work was supported by NIH/NINDS Grants U01NS098975 and U01NS123127, the Tianqiao & Chrissy Chen Brain–Machine Interface Center, and the Boswell Foundation. We thank S. Shimojo for valuable suggestions, V. Bougou and A. Betancourt Vera for providing code for data processing, J. Gamez for assistance with experiments and data processing, and T. Callier, B. Ruszala, S. Darcy, M. J. Thurston, S. K. Wandelt, and I. A. Rosenthal for helpful discussions.

## Author contributions

B.L. and R.A.A. conceived the project. B.L. and D.A.B. designed the tasks. B.L. collected the data. B.L., D.A.B., and L.S. analysed the data. B.L., D.A.B., L.S. and R.A.A. interpreted the data. C.Y.L. and B. Lee conducted the surgical procedures and provided clinical care. D.A.B. and K.P. led administration aspects of the clinical trial. B.L. and R.A.A. wrote the manuscript. All authors discussed and edited the manuscript. R.A.A. supervised the work.

## Additional information

Supplementary Information is available for this paper.

## Data availability

All data are available from the lead corresponding author upon reasonable request.

## Code availability

All code is available from the lead corresponding author upon reasonable request.

## Extended Data Figure legends

**Extended Data Fig. 1.**
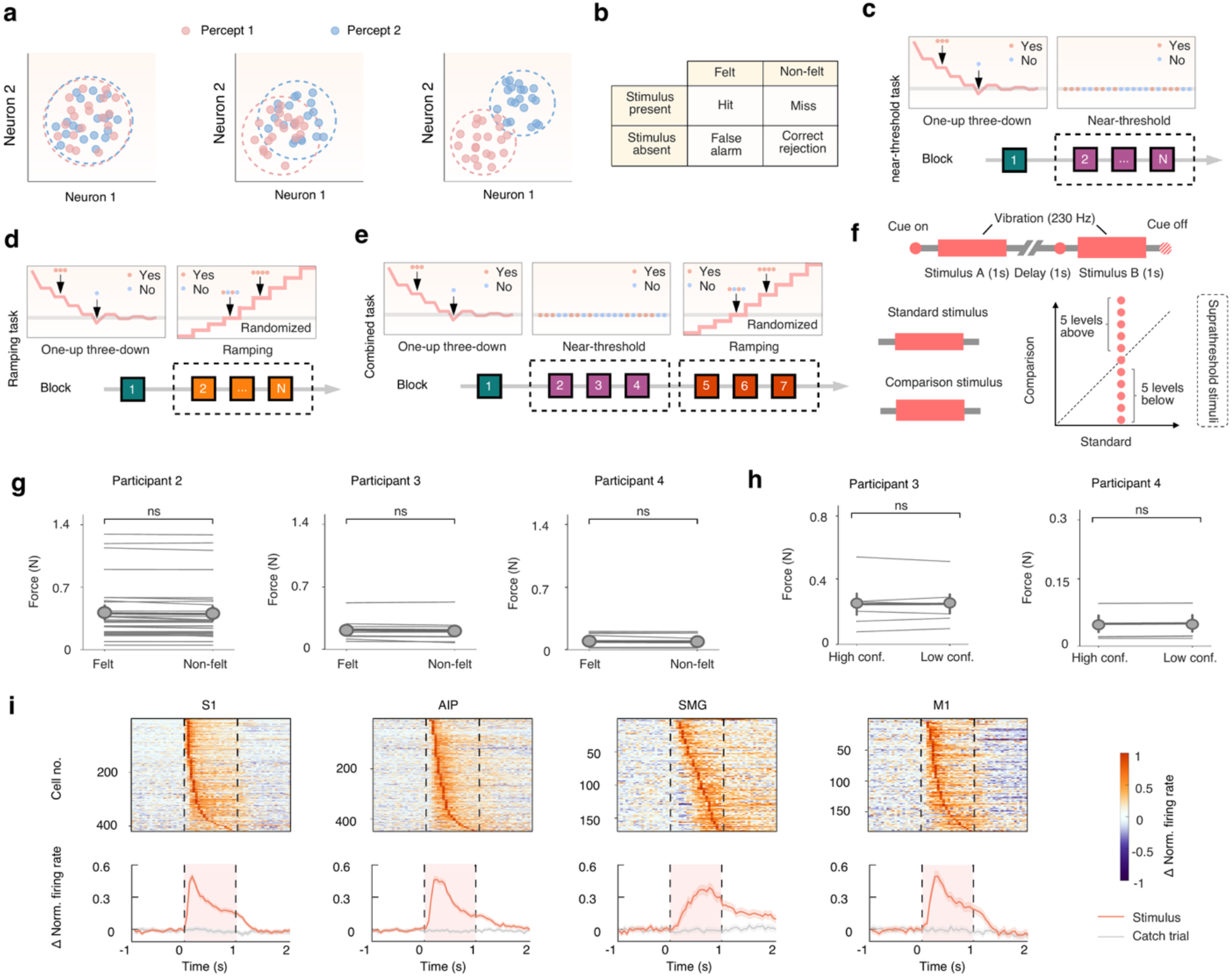
Task, behaviour, and distributed tactile responses across the grasp network. **a,** Three possible encoding schemes for subjective sensory perception in a brain area. Each dot denotes a sample neuron projected into a two-dimensional latent neural space. Colours denote perceptual outcomes. (Left) Neural activity patterns of the two percepts are largely overlapping, showing no discernible separation in latent space. (Middle) Neural populations exhibit partial separation between percepts, indicating limited discriminability. (Right) The two percepts form clearly distinct clusters in latent space, suggesting robust and separable encoding of subjective sensory perceptions. **b,** Outcome categories in the near-threshold detection task. Trials can be classified as hits or misses when the stimulus is present, and as false alarms or correct rejections during stimulus-absent catch trials, based on felt versus non-felt behavioural reports. **c**, Schematic of the near-threshold task procedure. Each session began with a one-up three-down staircase to estimate the participant’s vibrotactile detection threshold, followed by several near-threshold blocks in which the same amplitude was repeatedly presented to elicit variable felt and non-felt reports. **d**, Schematic of the ramping task procedure. After threshold estimation, several ramping blocks were performed in which vibrotactile stimuli spanning multiple amplitudes were randomly presented to characterize amplitude-dependent neural responses. **e**, Schematic of sessions combining near-threshold and ramping tasks. Threshold estimation was followed by several near-threshold blocks and ramping blocks to examine perceptual-state and amplitude coding in the same neurons. **f,** Schematic of the vibrotactile discrimination task. A 1-s, 230-Hz standard stimulus was presented first, followed after a 1-s delay by a 1-s comparison stimulus. The two intervals contained a fixed supra-threshold standard stimulus and a comparison stimulus. The comparison amplitude was randomly selected from 11 levels spanning five levels below, one level equal to, and five levels above the standard. After the cue disappeared, participants reported whether the comparison stimulus was stronger or weaker than the standard. **g**, Force measured by the accelerometer for felt and non-felt trials in the near-threshold task for each participant. Forces do not differ between felt and non-felt trials, indicating that subjective reports are not explained by input amplitude differences. **h**, Force measured by the accelerometer for high- and low-confidence trials in Participants 3 and 4. Forces do not differ between confidence levels, indicating that confidence ratings are not driven by physical amplitude. **i,** Normalized firing rate changes for stimulus trials and catch trials across the cortical grasp network. The upper panel shows latency-sorted individual cells across the three participants, and the lower panel shows the average across cells for each area. Stimulus trials are taken from the +100% condition (highest amplitude) in the ramping task, and the tactile modulated cells are shown. The results indicate that tactile stimulation elicits widespread activation across the grasp network.

**Extended Data Fig. 2.**
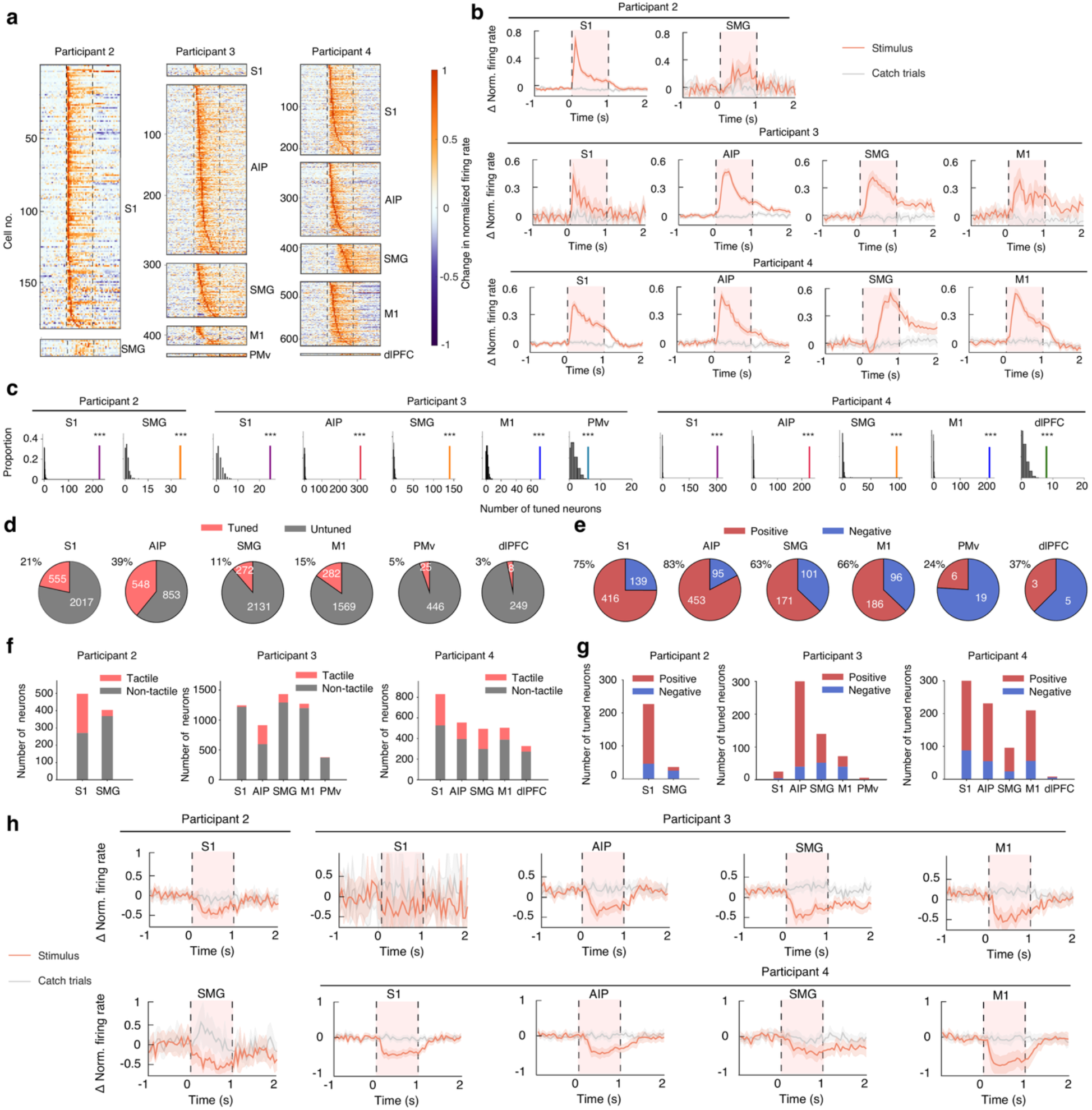
Neural profiles of tactile evoked activity in the grasp network. **a,** Normalized firing rate changes for supra-threshold (+100%) stimulus trials across recorded areas for each participant. This panel expands Extended Data Fig. 1i by showing participant-level excitatory responses. Dashed lines indicate the stimulus period. Stimulus trials are taken from the highest-amplitude (+100%) condition of the ramping task. Across participants, supra-threshold vibrotactile stimulation elicits robust, distributed responses throughout the grasp network, with inter-individual variability in amplitude and timing. **b,** Mean firing rate change for neurons classified as excitatory in a. Neurons are labelled as excitation response when their mean firing rate change during the stimulus window (0–1 s) exceeded 0. On average, excitation response shows prominent increases in firing across cortical areas. **c,** Observed number of tactile-tuned neurons compared with a shuffled distribution for each recorded area in the grasp network. The number of tactile-tuned neurons is significantly greater than expected by chance. **d,** Proportion of neurons tuned to vibrotactile stimulation among all recorded neurons, pooled across participants. Each pie chart shows the fraction of tuned (red) versus untuned (grey) neurons in each cortical region, highlighting that a substantial minority of neurons in every area encodes tactile information. **e,** Proportion of tactile-tuned neurons showing excitation (positive modulation; red) or inhibition (negative modulation; blue) responses across cortical regions, pooled across participants. Percentages indicate the fraction of tuned neurons showing positive modulation in each area, highlighting that most tuned neurons are positively modulated. **f,** Number of tactile-tuned neurons among all recorded neurons. A minority of neurons in each area are tuned to vibrotactile stimulation. For Participant 3, because most vibrotactile stimuli fell outside the receptive field of the S1 array, few S1 neurons were tactile tuned in the ramping task. S1 in Participant 3 was therefore not included in subsequent amplitude-sensitivity analyses. **g,** Number of excitation and inhibition responses among all tactile-tuned neurons. Most areas show a predominance of excitation responses over inhibition responses. **h,** Mean firing rate change for neurons classified as showing inhibition responses, defined as neurons whose mean firing rate change during the stimulus window (0–1 s) is below 0. This subset of the neuronal population in each area shows a clear decrease in firing after vibrotactile stimulation.

**Extended Data Fig. 3.**
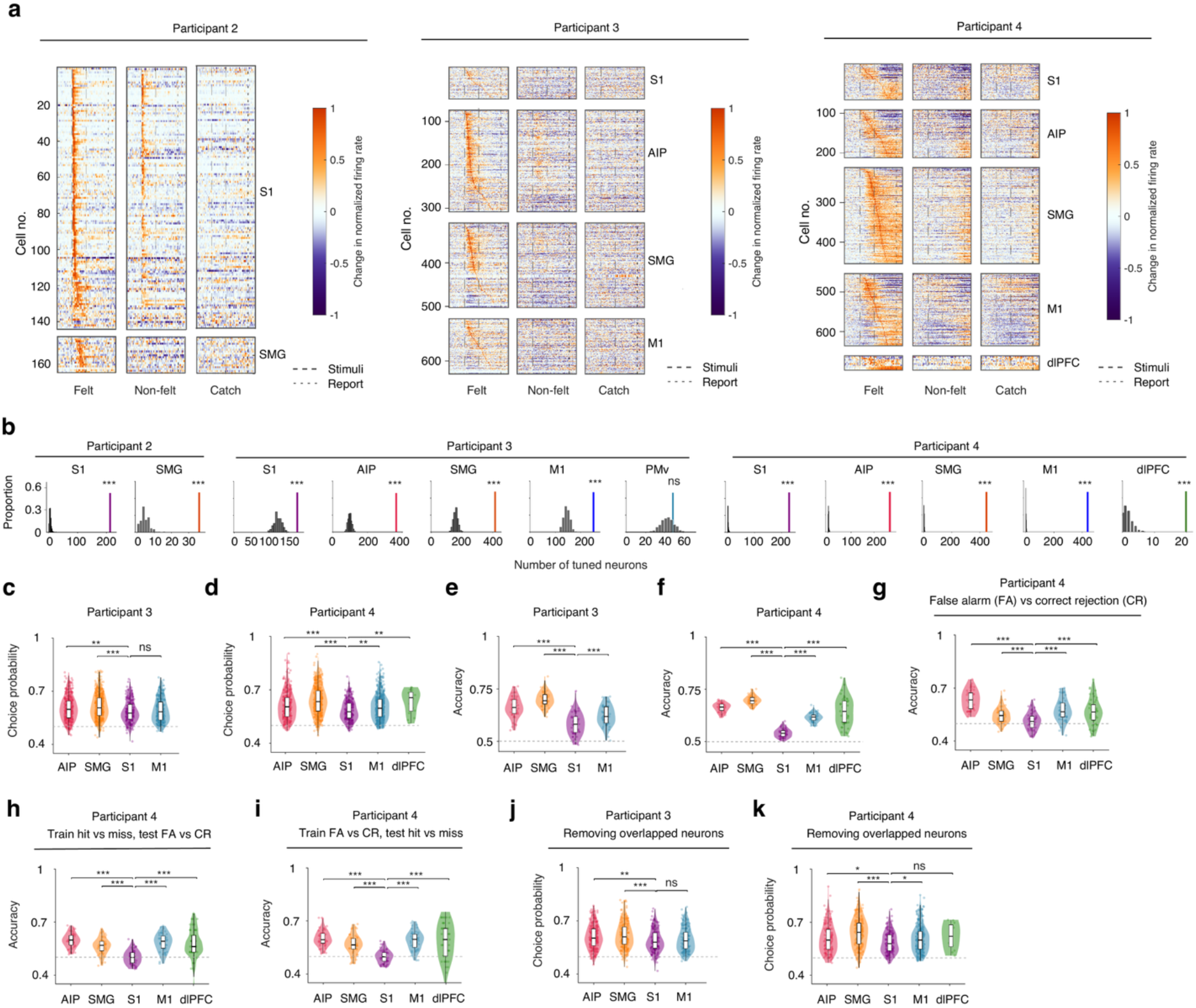
Neural responses during near-threshold stimulation. **a**, Expanded version of Fig. 1i showing normalized firing rate changes for felt and non-felt trials in each recorded area for individual participants, with neurons sorted by latency. Subjective near-threshold perception evokes prominent activation in PPC and S1, as well as M1, with inter-individual variability. In Participant 2, many S1 neurons also respond during non-felt trials, indicating encoding of the physical stimulus in addition to perceptual choice. Although only neurons with excitation responses above baseline are shown, some also exhibited sustained firing-rate suppression below baseline after stimulus offset. **b,** Number of neurons modulated near threshold across recorded areas compared with shuffled distributions. In the recorded areas except PMv, the number of near-threshold modulated neurons is significantly greater than expected by chance, revealing a distributed representation of subjective tactile perception across the grasp network. **c**,**d,** Expanded version of Fig. 2f,g showing choice probability for distinguishing hit and miss trials across cortical areas. S1 showed significantly lower choice probability than AIP and SMG in both participants and than M1 and dlPFC in Participant 4 (Benjamini– Hochberg FDR-adjusted *P* < 0.05). The difference between S1 and M1 in Participant 3 was not significant. PMv in Participant 3 was not shown in c,e,j because its number of near-threshold-modulated neurons did not exceed chance in b. **e**,**f,** Expanded version of Fig. 2f–g showing neuron-dropping decoding accuracy for distinguishing hit and miss trials across cortical areas, with neuron count matched across areas. S1 showed significantly lower decoding accuracy than the other cortical areas, including M1 in both participants, indicating weaker population-level encoding of subjective tactile perception. **g**–**i**, Population decoding and cross-decoding of perceptual outcome in Participant 4. Violin plots summarize decoding accuracy across brain areas for false alarm (FA) versus correct rejection (CR) trials (g), training on hit versus miss and testing on FA versus CR (h), and training on FA versus CR and testing on hit versus miss (i). All analyses used matched decoding parameters across areas. Dots indicate individual decoding repeats, the dashed line marks chance level, and statistically significant pairwise comparisons are indicated. Higher decoding accuracy in PPC and other non-S1 areas than in S1, across stimulus-present and stimulus-absent conditions, suggests that these regions encode a more abstract perceptual representation, whereas S1 carries a less generalizable sensory code. **j**,**k**, Choice probability for Participant 3 (j) and Participant 4 (k) after removing neurons that overlapped across recording sessions. Other analysis and plotting settings are as in c,d. PPC areas retained higher choice-related information than S1 after excluding these putatively repeated neurons, indicating that the PPC advantage was not driven by cross-session neuron overlap.

**Extended Data Fig. 4.**
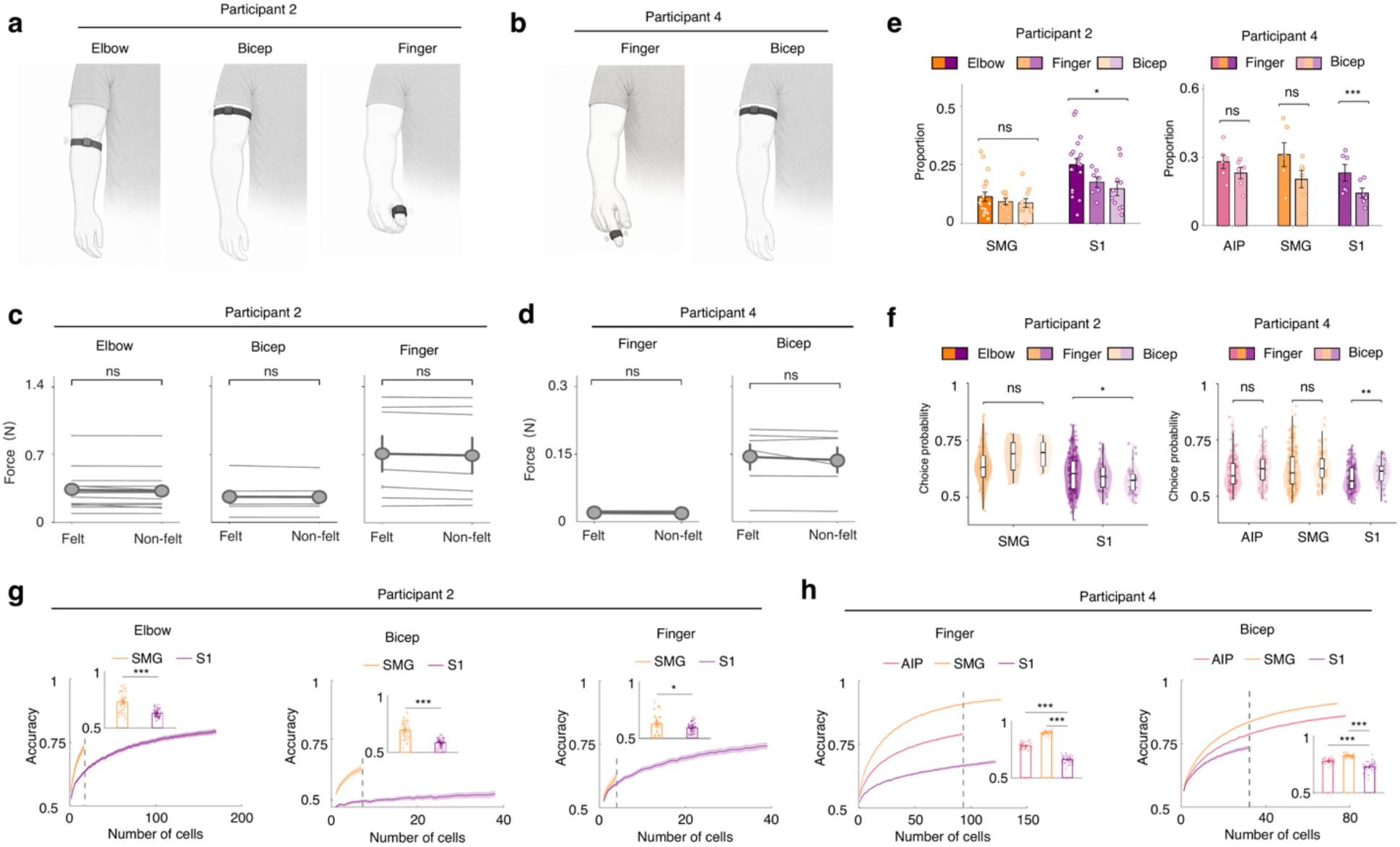
Vibrotactile stimulation location and its effects on neuronal correlates of subjective tactile awareness. **a**,**b,** Schematic of vibrotactile stimulation location in two participants (a, Participant 2; b, Participant 4). For Participant 2, vibrotactile stimulation was delivered at three sites with preserved sensation: the bicep (anteromedial), elbow (anterior cubital fossa), and finger (lateral base of the thumb near the thenar eminence). For Participant 4, vibrotactile stimulation was tested at two sites with intact sensation: the bicep (anteromedial) and the finger (tip of the index finger). The stimulation locations were tested in separate sessions. **c**,**d**, Force measured by the accelerometer for felt versus non-felt near-threshold trials, shown per participant (c, Participant 2; d, Participant 4) and separated by vibrotactile location. Within each location, forces are comparable between felt and non-felt trials, although perceptual thresholds differ across locations, consistent with differences in skin sensitivity or injury level. **e**, Session-wise proportion of modulated near-threshold neurons for each vibrotactile location (elbow, finger and bicep for Participant 2; finger and bicep for Participant 4) in PPC and S1. In S1, the fraction of near-threshold sensitive neurons differs across somatotopic locations and between participants, whereas differences in PPC are less pronounced. **f,** Choice probability for distinguishing felt from non-felt trials for each vibrotactile location. In S1, choice probability varies significantly across locations and participants, consistent with somatotopic differences, whereas the difference in PPC choice probability is not statistically significant. Together with the results in e, these findings suggest less divergence across locations and weaker somatotopy effects in PPC. **g**,**h,** Neuron-dropping curves for decoding felt versus non-felt trials for each vibrotactile location, extending the analysis in Fig. 2e–g. Insets show decoding accuracy when the number of neurons is matched, as indicated by the dashed line. Across locations, PPC areas consistently achieve higher decoding accuracy than S1.

**Extended Data Fig. 5.**
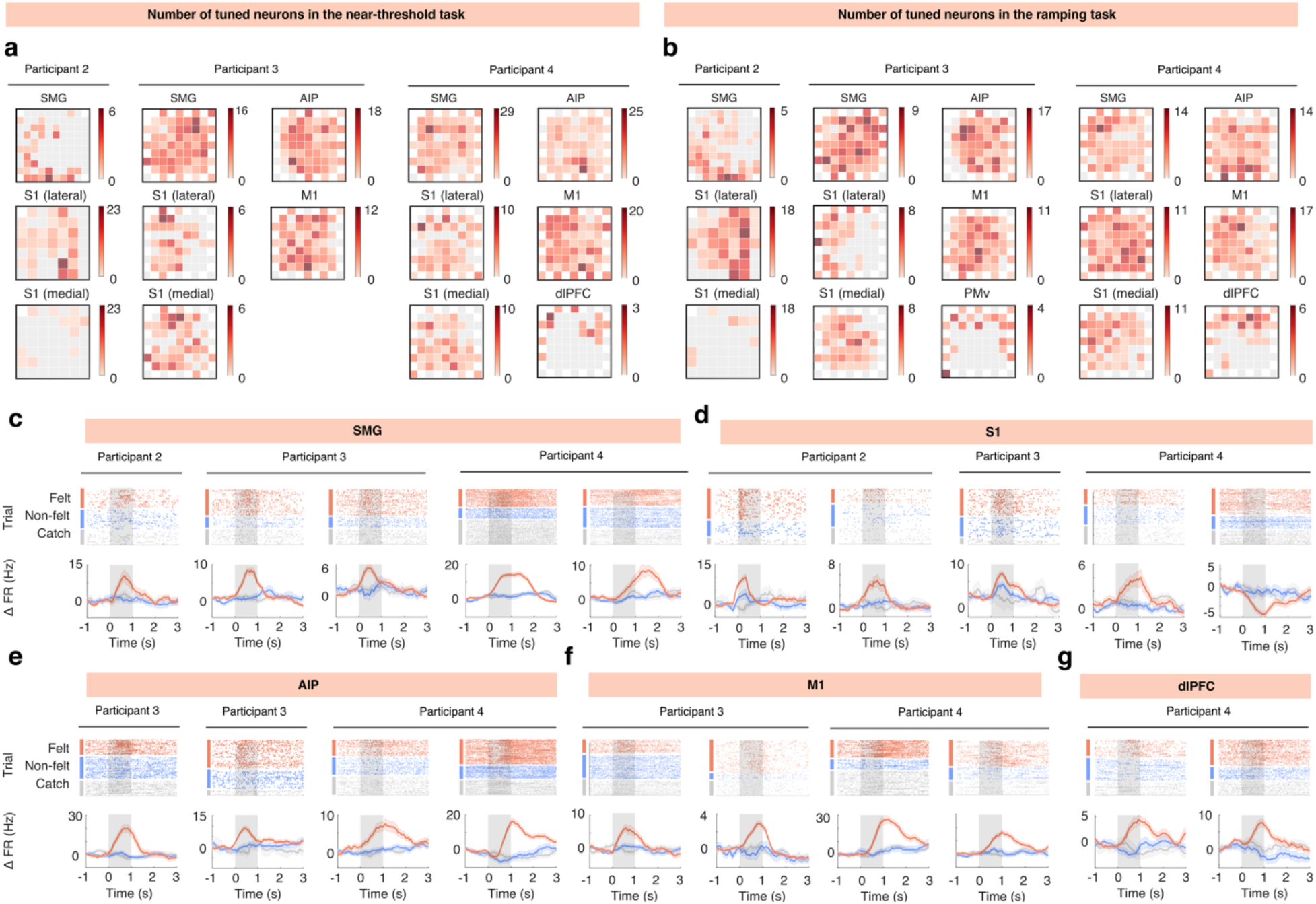
Spatial and temporal profiles of tuned neurons across recording arrays and participants. **a,** Topographic maps of near-threshold modulated neurons in recorded arrays during the near-threshold task. Multiple electrodes contribute across sessions, arguing against repeated sampling from a single site and showing a broad spatial distribution. **b,** Topographic maps of modulated neurons during the ramping task. For both a and b, neurons are pooled across sessions and assigned to their recording channel. Colour indicates the number of modulated neurons per channel. **c**–**g,** Response profiles of neurons modulated near threshold, grouped by cortical area and participant. Raster plots show trial-by-trial firing rate changes for felt, non-felt and catch trials, and traces below show the corresponding average firing dynamics. Felt trials showed stronger responses, with variable timing and magnitude across participants.

**Extended Data Fig. 6.**
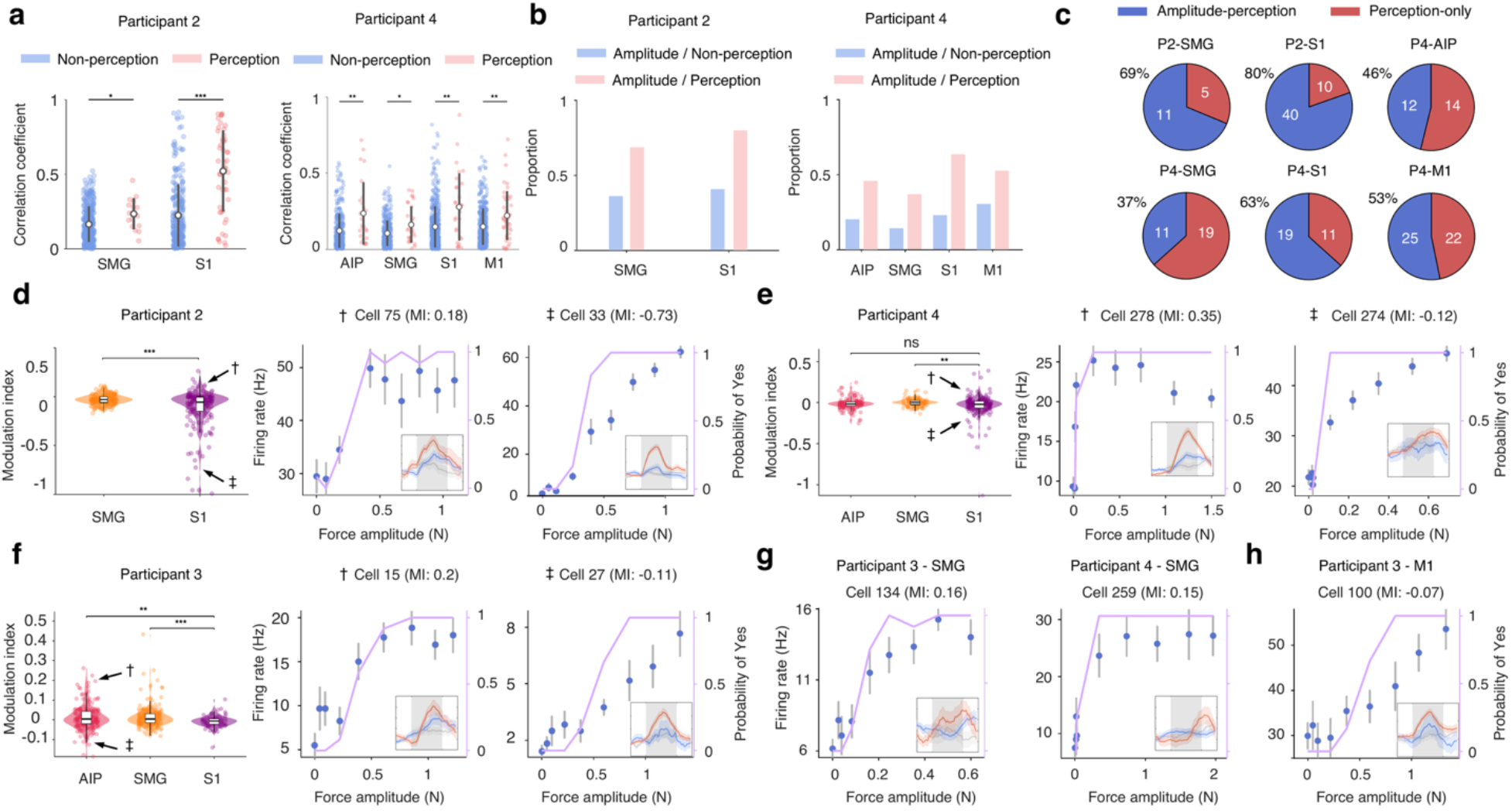
Perceptual-state and amplitude coding using combined near-threshold and ramping task. **a,** Correlation between firing rate and stimulus amplitude during the ramping task for neurons with and without near-threshold modulation. Neurons with near-threshold modulation showed stronger amplitude correlations, suggesting that there are not two classes of neurons and the response properties lie along a continuum. **b,** Proportion of amplitude-modulated neurons among neurons with and without near-threshold modulation. Near-threshold modulated neurons were more likely to be amplitude tuned, supporting a continuum. **c,** Overlap between perceptual-state modulation and amplitude modulation in PPC, S1 and M1. Pie charts show neurons with both amplitude and perceptual modulation versus perception-only modulation, revealing both mixed-selective and perception-predominant response profiles. Participant 3 was not included in the comparisons in a–c because the stimulation site lay outside the receptive fields of the recorded S1 neurons. **d**–**f**, Modulation indices and example neuronal response profiles for Participants 2 (d), 4 (e) and 3 (f). Left, GLM-based modulation indices for neurons with near-threshold or amplitude modulation. Lower values indicate relatively stronger amplitude modulation, whereas higher values indicate relatively stronger near-threshold modulation. Middle and right, example S1 (d,e) and AIP (f) neurons showing saturating sigmoidal responses (middle, †) and graded responses across stimulus amplitudes (right, ‡). Arrows and matching symbols identify the example neurons in the modulation index distributions. Blue points show firing rates during the ramping task, and purple lines indicate behavioural detection probability (right axes). Insets show felt (red) and non-felt (blue) responses during the near-threshold task. Modulation indices indicated a stronger relative preference for amplitude coding in S1 than in SMG, with S1 and AIP neurons exhibiting both graded ramping responses and saturating sigmoidal responses as stimulus amplitude increased. **g**,**h**, Examples of saturating sigmoidal responses in SMG (g; Participant 3, left; Participant 4, right) and a graded ramping response in M1 (h; Participant 3). Graded ramping and saturating sigmoidal response profiles were also observed beyond S1 and AIP, although less commonly.

**Extended Data Fig. 7.**
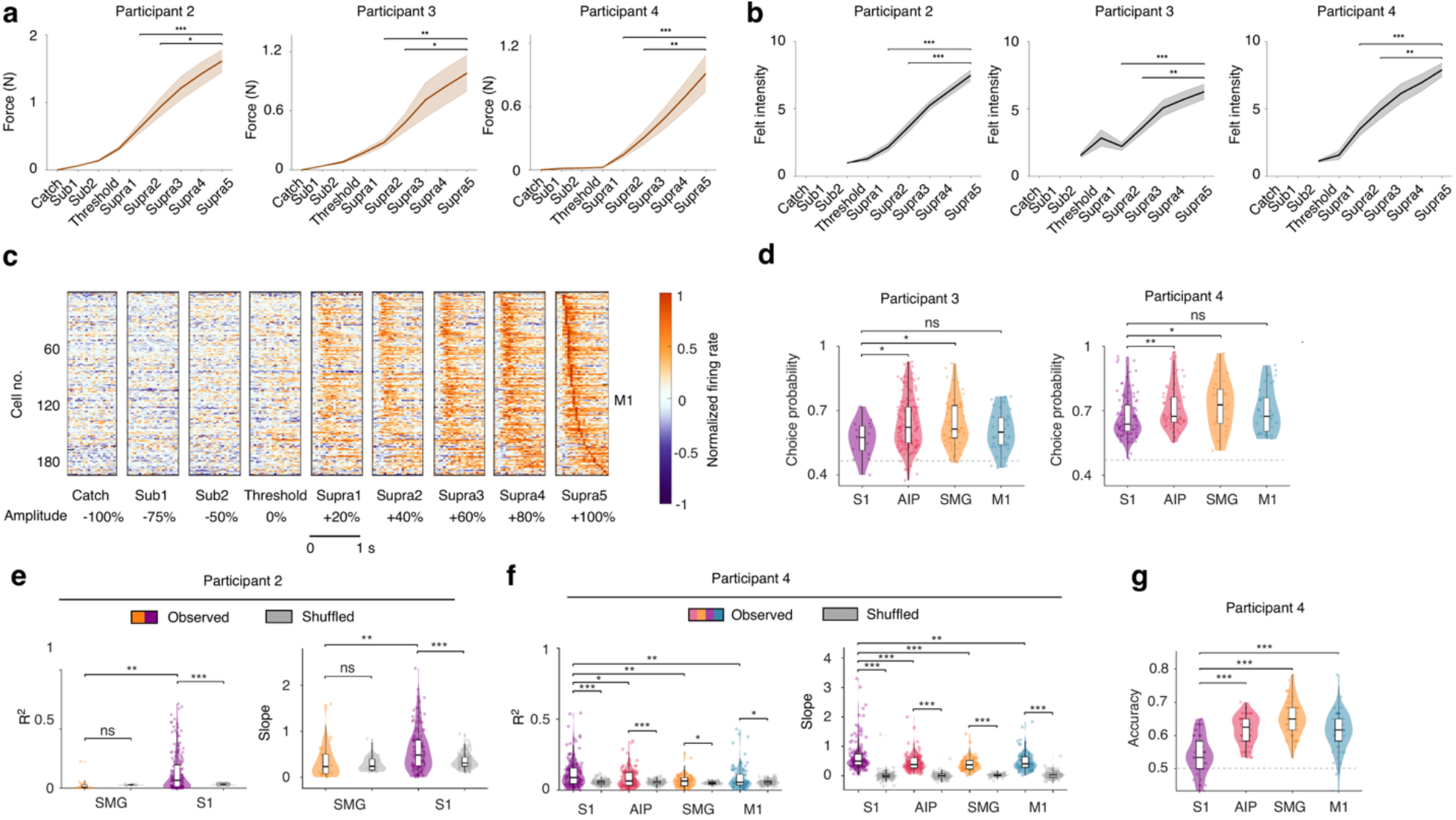
Decoupling stimulus amplitude and perceptual reports in the ramping task. **a,** Measured force increased with commanded amplitude across ramping conditions, confirming graded physical stimulation. **b,** Behavioural felt intensity increased with stimulus amplitude, showing that the ramping task sampled a broad perceptual intensity range. **c,** Expanded version of Fig. 3a with normalized firing rate changes in M1 across ramping amplitudes, revealing amplitude-dependent modulation in motor cortex. **d,** Choice probability for distinguishing felt and non-felt responses using near-threshold trials from the ramping task. Consistent with Fig. 2f,g, PPC exhibits higher choice probability than S1, supporting a greater contribution of PPC to perceptual choice. **e,** Subliminal analysis of firing rate–force relationships in non-felt trials. R² and absolute slope values were compared with shuffled distributions, showing significant subliminal amplitude modulation in S1 but not PPC, and indicating stronger amplitude encoding in S1 in the absence of conscious perception. **f,** Supraliminal analysis of firing rate–force relationships within each felt-intensity level. R² and slope values were compared with shuffled distributions, showing significant amplitude modulation in both PPC and S1, with stronger amplitude encoding in S1. **g,** Decoding high-versus low-felt intensity trials after fixing physical amplitude in the ramping task. This analysis tested perceptual-intensity representation by controlling for stimulus strength and showed stronger perceived-intensity signals in PPC than S1.

**Extended Data Fig. 8.**
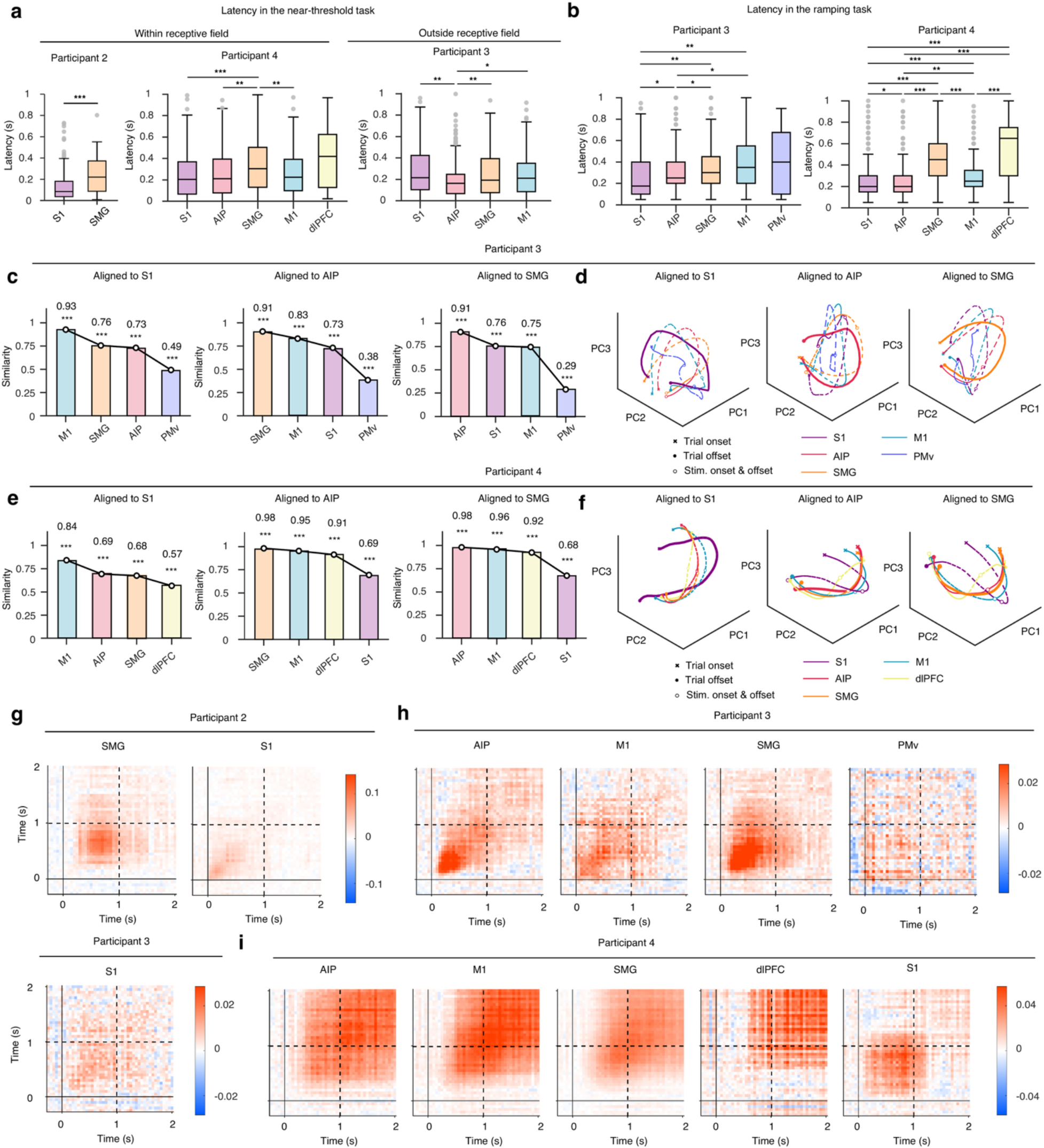
Temporal hierarchy and population-level structure of tactile perceptual representations. **a**,**b**, Response latency across participants and brain areas in the near-threshold and ramping tasks. Boxplots summarize neuronal response latency for Participants 2–4 in the near-threshold task (a) and Participants 3–4 in the ramping task (b, multi-region extension of Fig. 3l). Latency was defined for each neuron as the first time bin with a significant firing-rate difference for felt versus non-felt trials in a and +100% versus catch trials in b. Statistically significant pairwise comparisons are indicated. Across both the near-threshold task within the receptive field (Participants 2 and 4) and the ramping task (Participants 2, 3 and 4; results for Participant 2 are shown in Fig. 3l), response latency showed a trend toward an S1 < AIP < SMG pattern, suggesting hierarchical processing from early somatosensory to higher-order parietal areas. **c**–**f**, Cross-area alignment of low-dimensional neural trajectories in Participants 3 and 4. c,e, Procrustes similarity between each target area and three reference areas, S1, AIP and SMG, for Participant 3 (c) and Participant 4 (e). Neural activity from perception-sensitive neurons was averaged for felt trials in the near-threshold task, projected onto the top three principal components, and aligned to each reference area using Procrustes transformation. Bars show the similarity between the aligned target-area trajectory and the reference-area trajectory. Numbers indicate similarity values. Statistical significance was assessed using 1,000 permutation tests; \*\*\**P* < 0.001. d,f, Three-dimensional PCA trajectories after Procrustes alignment for Participant 3 (d) and Participant 4 (f). Each plot shows the trajectories of recorded brain areas after alignment to S1, AIP or SMG (left to right). Colours indicate brain areas. Bold lines denote the reference area used for alignment. Crosses and filled circles denote trial onset and offset, respectively; open circles denote stimulus onset and offset. The results suggest that M1 and S1 share more similar neural manifolds, whereas the two PPC regions, AIP and SMG, exhibit a more closely aligned manifold structure. **g**–**i**, Cross-time representational similarity analysis (RSA) in the near-threshold task. Pseudo-population RSA maps summarize the temporal structure of felt versus non-felt neural activity across brain areas in three participants. Activity from perception-sensitive neurons was used to construct area-specific pseudo-populations with balanced felt and non-felt trials. Colour values represent a correlation-based RSA contrast for each pair of time bins, computed as the mean similarity among trials with the same perceptual outcome minus the mean similarity between felt and non-felt trials across pseudo-population activity vectors. Solid lines mark stimulus onset, and dashed lines mark stimulus offset. Across participants, sustained RSA structure in PPC (SMG and AIP) suggests a role in maintaining tactile perceptual representations beyond the temporally restricted S1 response.

**Extended Data Fig. 9.**
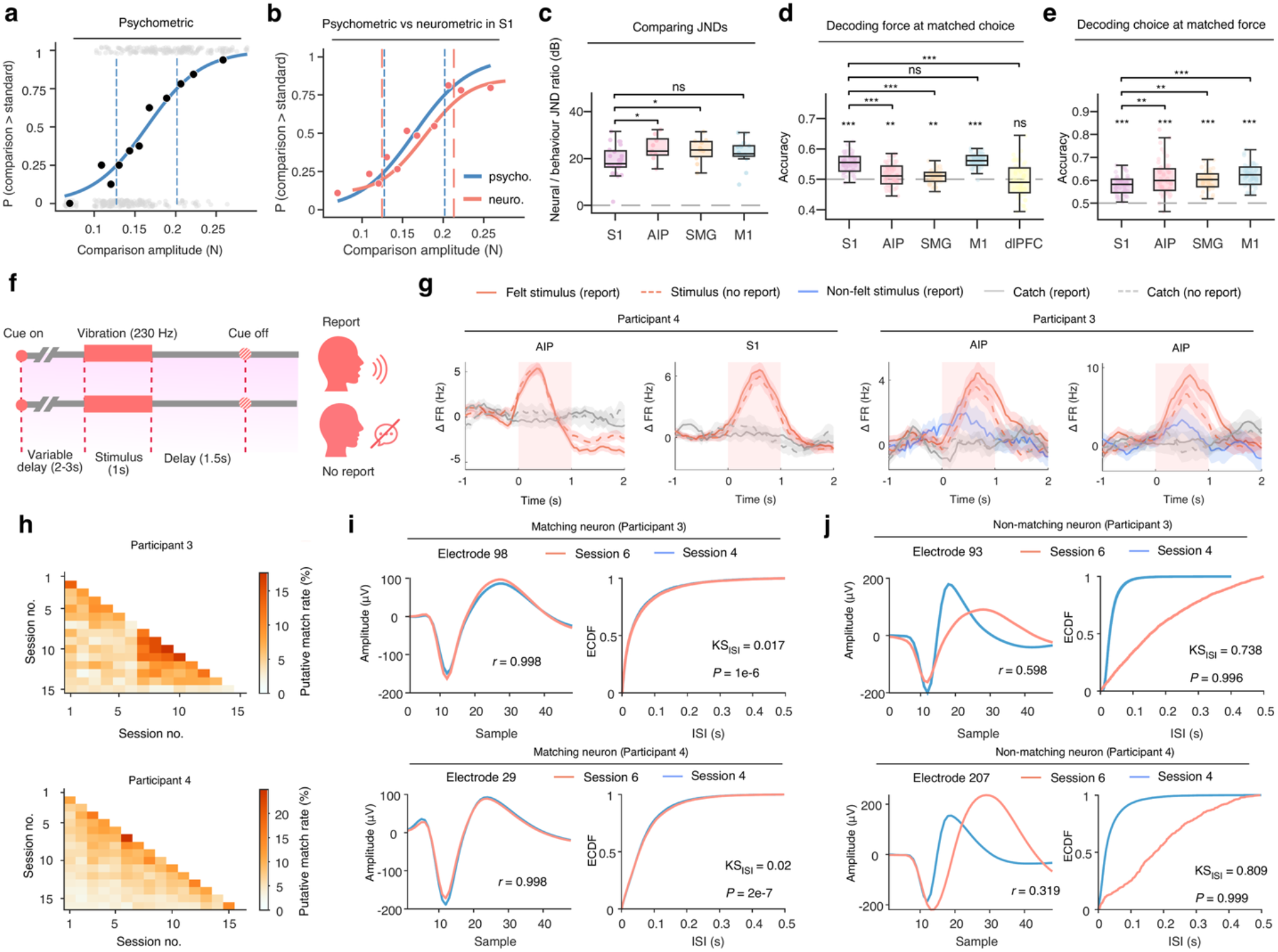
Control analyses of vibrotactile discrimination task, no-report task and cross-session neuronal stability. **a**, Example psychometric function for the vibrotactile discrimination task. The blue line shows a logistic fit to trial level choices using maximum likelihood estimation. Grey dots indicate individual choices, black circles indicate response proportions, and dashed lines mark the 25% and 75% points used to calculate the just noticeable difference (JND). **b**, Psychometric and neurometric curves for a representative S1 neuron. Red dots show neural choice proportions. Blue and red dashed lines mark the psychometric and neurometric JND bounds, respectively. Their similar spacing indicates that this S1 neuron tracks physical sensitivity to differences in force. **c**, Comparison of neural-to-behavioural JND ratios across cortical areas. For each neuron, the neurometric JND was divided by the corresponding behavioural JND, converted to decibels, and grouped by cortical area. S1 showed lower ratios than AIP and SMG but not M1, indicating closer correspondence between neural and behavioural sensitivity to force differences. **d**, High and low force trials within the psychometric JND range were sampled to match behavioural choice, and equal numbers of neurons were used across areas. Neural population activity was used to classify high and low force levels. Dots show decoding repeats, and the dashed line indicates chance level. Markers above individual areas indicate significance versus chance, whereas brackets indicate pairwise comparisons. S1 and M1 achieved higher accuracy than PPC, indicating stronger encoding of physical force independent of choice. **e**, Decoding behavioural choice at matched force. Trials were sampled to match physical force distributions across choices, and equal numbers of neurons were used across areas. Neural population activity classified whether the comparison was judged stronger or weaker than the standard. AIP, SMG and M1 outperformed S1, indicating stronger choice related encoding independent of force. **f**, No-report control task. To control for neural activity associated with explicit perceptual reporting, participants were not required to report their perception in some blocks of the near-threshold task. Trial structure and stimulation were the same as in the standard task, except that participants reported felt or non-felt at cue offset only in report blocks. Report and no-report blocks, both including vibration and catch trials, were interleaved. **g**, Neural responses during report and no-report blocks. Mean baseline-corrected firing-rate changes are shown for felt stimulus and non-felt stimulus trials in report blocks, stimulus trials in no-report blocks, and catch trials in both report and no-report blocks across representative PPC and S1 neurons in Participants 3 and 4. Solid and dashed lines indicate report and no-report conditions, respectively. Similar responses across report and no-report conditions suggest that these neural responses persisted in the absence of overt reporting. **h**, Cross-session putative neuron matching rates for Participants 3 and 4. Heatmaps show the percentage of neurons identified as putative matches between each pair of recording sessions. Matching rates were generally low across most session pairs (Participant 3: median, 5.08%; range, 1.42–17.66%; Participant 4: median, 6.29%; range, 2.56–24.96%), indicating limited overlap in putatively stable neurons across recording days. **i**, Representative putatively matching neurons from Participants 3 and 4. For each example, average spike waveforms (left) and inter-spike interval (ISI) empirical cumulative distribution functions (ECDFs; right) are compared between two sessions. Waveform similarity is indicated by the Pearson correlation coefficient (*r*), and differences between ISI distributions are quantified by the Kolmogorov–Smirnov statistic (KS_ISI_); *P* denotes the joint matching *P* value calculated using Fisher’s method. Despite low cross-session matching rates, neurons with highly similar spike waveforms and ISI distributions were identified. **j**, Representative non-matching neurons from Participants 3 and 4, shown using the same waveform and ISI comparisons as in i. Non-matching neurons exhibited lower waveform correlations and larger differences between ISI distributions than the putatively matching examples.

**Extended Data Fig. 10.**
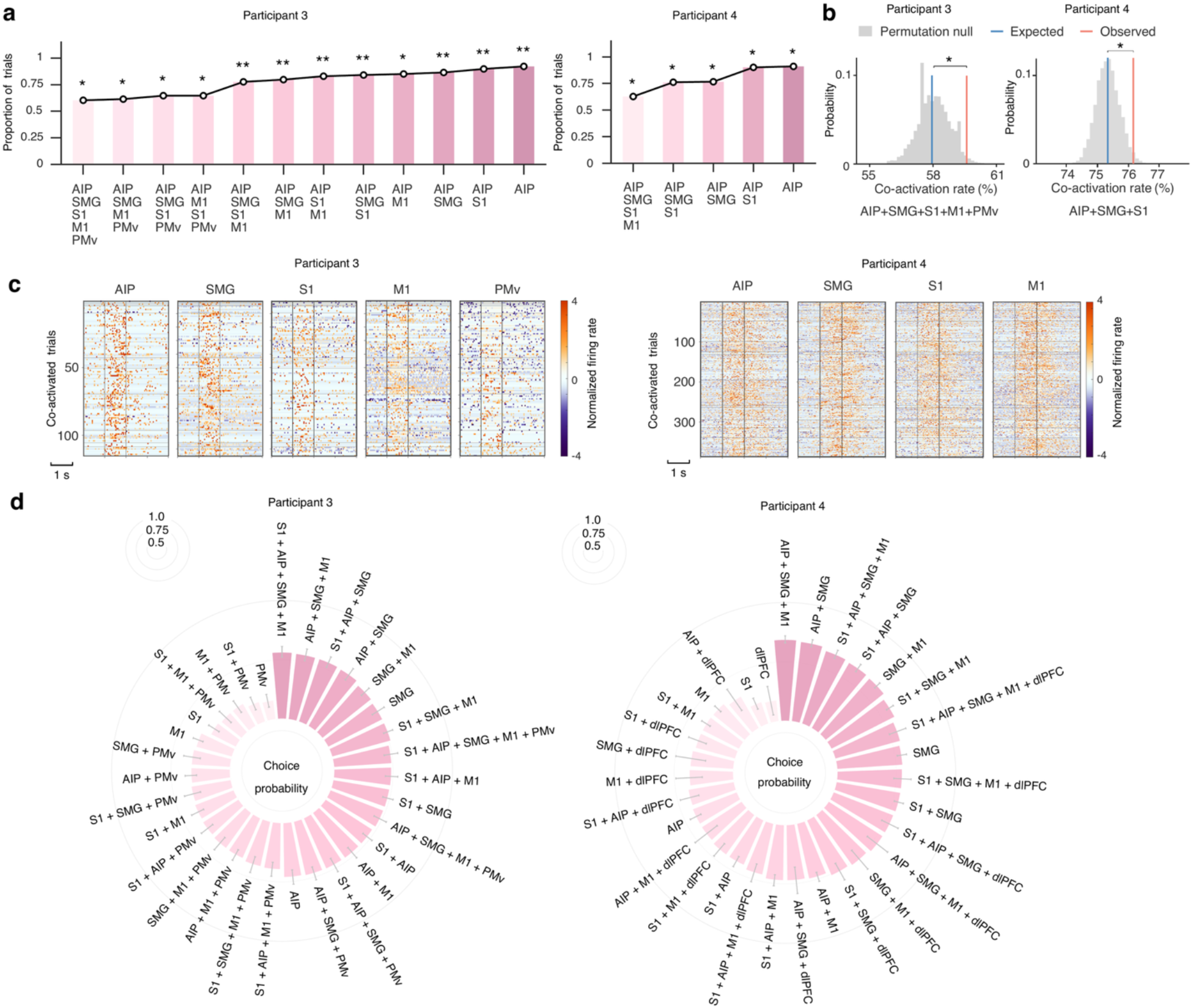
Simultaneous activation across cortical areas is associated with subjective tactile perception. **a**, Proportion of felt stimulus trials showing significant activation in single cortical areas or significant simultaneous activation across multiple cortical areas, i.e., *P* (Activation Felt), in Participants 3 and 4. Significance was assessed against a null distribution generated by shuffling trial labels within each session. Only significant combinations are shown. These significant activation patterns across different area combinations indicate that conscious tactile perception can be supported by different subsets of cortical areas at the single-trial level. **b**, Example cortical area combinations showing significantly higher observed co-activation than expected under independence during felt trials in both participants. Expected co-activation probability was defined as the product of the individual area activation probabilities, ∏*_i_ P_i_* (Activation ∣ Felt) . The observed co-activation rate was significantly higher than the product of the individual area activation probabilities in both participants, supporting coordinated multi-area activation during conscious tactile perception. **c**, Single-trial firing-rate rasters for felt trials with simultaneous perception-sensitive activation across example combinations of cortical areas. Each row represents the same simultaneous felt trial across area panels. Vertical lines indicate stimulus onset and offset, and horizontal lines separate recording sessions. These single-trial responses demonstrate concurrent neuronal activation across distributed cortical regions during individual felt trials. **d**, Choice probability for discriminating felt from non-felt trials based on single-neuron activity in individual cortical areas and all possible area combinations. Bar length indicates the mean choice probability across sessions, with error bars showing s.e.m.; combinations are ordered by choice probability. Choice probability was generally higher for multi-area combinations than for individual areas, indicating that distributed cortical activity across multiple regions provides greater information about perceptual outcome than activity within a single region alone.

## Methods

### Clinical trial and participants

Data were collected from three male participants (Participants 2, 3, and 4) with chronic tetraplegia. Before enrolment, prospective participants completed a detailed consent process during which they were informed that participation was not expected to provide any direct benefit. All participants provided written informed consent and were enrolled in an ongoing clinical trial (NCT01964261) conducted under an Investigational Device Exemption granted by the US Food and Drug Administration. Further information about the parent trial is publicly available on https://clinicaltrials.gov/study/NCT01964261. Participant numbering was retained from the clinical trial labels (S2, S3, and S4), with participants referred to as Participants 2, 3, and 4 to avoid confusion with cortical area names. The study protocol was approved by the Institutional Review Boards of California Institute of Technology, the University of Southern California, and Rancho Los Amigos National Rehabilitation Center. The main data presented span a period of two years as part of the longitudinal clinical trial. Recording sessions were typically conducted on separate days, each lasting 2 to 3 hours, including preparation and breaks.

Participant 2 had a C5-level spinal cord injury, 8.5 years post-injury and 7 years post-implant at the time of the first session. Residual sensations were present in the shoulder, bicep, elbow, and lateral thumb. Participant 3 had a C5-level spinal cord injury, 3.5 years post-injury and 1 year post-implant at the first session, exhibiting residual sensation in the upper arm but largely absent or unnatural percepts across the hand and forearm. Participant 4 sustained a C7-level spinal cord injury 50 years earlier, was 3 months post-implant at the first session, and showed intact sensation in the anterior forearm and most fingertips (except the pinky) with residual sensation in the posterior arm. A subset of participants was used for some analyses based on the availability of recordings for the specific experimental paradigm and cortical areas examined.

### Implant and single-neuron recordings

All participants completed preoperative anatomical and functional MRI to localize grasp-related cortical areas for guiding surgical strategy and array placement.

Participant 2 received four implanted Utah microelectrode arrays targeting supramarginal gyrus (SMG), primary somatosensory cortex (S1), and ventral premotor cortex (PMv). The arrays in SMG and PMv were platinum-tipped NeuroPort arrays (Blackrock Microsystems, Salt Lake City, UT) with 96 channels each, whereas the two S1 arrays (medial and lateral) were sputtered iridium oxide film (SIROF)-tipped microelectrode arrays with 48 channels each. Due to substantial signal degradation in PMv recordings during years 7–9 post-implant, data from PMv in Participant 2 were not used for analysis. Participants 3 and 4 each received six Utah arrays implanted in sensorimotor, parietal and frontal regions, with 64 channels per array. In both participants, two arrays targeted medial and lateral subregions of S1. For Participant 3, the additional arrays targeted SMG, the anterior intraparietal area (AIP), PMv, and the primary motor cortex (M1). For Participant 4, the additional targets included SMG, AIP, M1, and the dorsolateral prefrontal cortex (dlPFC). In Participants 3 and 4, the AIP arrays were placed on the medial gyrus at the intersection of the postcentral and intraparietal sulci, and the SMG arrays were placed in the inferior parietal lobule. In all participants, the S1 arrays were placed in the hand region (spanning the thumb to the little finger) of the postcentral gyrus. Each Utah microelectrode array had a footprint of 4 × 4 mm.

Broadband neural signals were recorded through three percutaneous connectors in each participant, yielding 288 channels for Participant 2 and 384 channels for Participants 3 and 4. For Participant 2, recordings were acquired using a Neural Signal Processor (Blackrock Microsystems, Salt Lake City, UT). For Participants 3 and 4, recordings were primarily acquired using Grapevine processors (Ripple Neuro, Millcreek, UT). The broadband neural signals were sampled at 30 kHz and stored for offline analysis.

For single-neuron analyses, task-modulated neurons were pooled across recording sessions within each participant and cortical area. The terms neuron, cell and unit are used interchangeably in the study to refer to spike-sorted units.

### Spike detection

Spike events were detected from broadband neural signal using a denoising and re-thresholding pipeline. Signals were band-pass filtered from 250 to 5,000 Hz with an eighth-order filter. Notch filters and a 60 Hz comb filter were applied when needed. For each channel, background noise was estimated based on a Gaussian noise model, and thresholds were set as negative multiples of the estimated noise level. The default threshold was set to −4.5 times the estimated noise level for Participants 3 and 4. For Participant 2, a less conservative threshold of −3.5 was used due to elevated noise.

### Spike sorting

Spike sorting was performed using a custom MATLAB-based framework, which included waveform alignment, outlier rejection, principal component analysis, and unsupervised clustering. The number of clusters was automatically estimated using gap or silhouette metrics. Clusters were evaluated based on signal-to-noise ratio and silhouette values, and assigned quality and unit type labels using pre-trained classifiers. Spike sorting was conducted separately for each session, and this pipeline enables fully automated processing that ensures reproducible outputs.

### Vibrotactile device

We developed a custom vibrotactile stimulation device by integrating a tactor motor (C3, Engineering Acoustics, Casselberry, FL) and a tri-axis accelerometer (LilyPad ADXL335, SparkFun Electronics, Niwot, CO, USA) into an elastic armband (Freedom Band, Fresno, CA). The tactor delivered 230 Hz vibrations lasting 1 second and was driven by a signal generator triggered from MATLAB. For Blackrock systems, the built-in waveform generator on the Neural Signal Processor was used. For the Ripple systems, stimulation was administered via an external signal generator (NI USB-6001, National Instruments, Austin, TX) connected to the recording computer. The generator output was amplified using an analog audio amplifier (Nobsound NS-01G PRO), which allowed manual adjustment of vibration amplitude. A custom-labelled dial was affixed to the amplifier to indicate relative gain settings.

To measure the actual vibratory output, the accelerometer was epoxied directly onto the surface of the tactor. It recorded acceleration along the horizontal, vertical, and depth directions, and was powered via digital outputs from the neural recording system. The three-axis acceleration signals, along with the analog signal driving the tactor, were recorded via analog input channels of the neural acquisition system. The analog stimulation signal served as the ground-truth trigger to determine the actual onset of each vibrotactile pulse.

### Vibrotactile trial structure

Each block began with on-screen instructions indicating the task type and experimental instruction. Individual trials began with the appearance of a central fixation dot on a blank screen. To preclude temporal expectation, the vibrotactile stimulus was delivered after a variable delay of 2 to 3 seconds and the vibration lasted for 1 second. Following a 1.5-second post-stimulus delay, the fixation dot disappeared, prompting participants to verbally provide a subjective report. The perceptual response of felt or non-felt, along with a confidence rating on a scale from 1 (lowest) to 10 (highest), was manually recorded by the experimenter. Participants were instructed to report only after the fixation cue had disappeared. To monitor false alarms, a proportion of trials ranging from 30 to 50 percent were designated as catch trials in which no stimulus was delivered. Participants were not informed about the presence or frequency of these trials.

### Vibrotactile stimulation location

To test whether neural correlates of subjective sensory perception were location-dependent, vibrotactile stimuli were applied to different sensate locations on the upper limbs of the participants in the near-threshold task. For Participant 2, three distinct locations were tested, the lateral base of the thumb near the thenar eminence, the anterior elbow region (cubital fossa), and the anteromedial region of the bicep. The elbow location corresponded to the somatotopic region represented by the implanted microelectrode array, where intracortical microstimulation (ICMS) reliably evoked tactile percepts^49,50^. For Participant 3, vibrotactile stimulation was delivered only to the bicep due to the absence of intact sensation on the forearm and hand. For Participant 4, two sites with preserved sensation were tested, i.e., the anteromedial biceps and the tip of the index finger. In both Participants 3 and 4, ICMS over S1 reliably evoked hand and finger sensations, indicating that the somatotopic representation of the hand was covered by the array. In the ramping task, vibrotactile stimulation was delivered to the elbow, bicep and finger for Participants 2, 3 and 4, respectively. These above locations were verified by preliminary mapping of tactile sensitivity across each participant’s arm and hand using Semmes–Weinstein monofilaments of varying forces to confirm that participants could detect touch at the selected sites.

### One-up three-down task

At the beginning of each session, we estimated the participant’s vibrotactile detection threshold using a one-up three-down staircase procedure^51,52^. In this adaptive protocol, stimulus amplitude decreased by a step size Δ*S_t_* after three consecutive “Yes” responses and increased by Δ*S_t_*_–1_/2 following a single “No” response. The staircase terminated after five reversals in direction, which was taken as convergence. The detection threshold was defined as the average amplitude over the final ten steps. To accelerate convergence, we manually adjusted the analog amplifier and the task began at a low-intensity, supra-threshold amplitude *S*_0_ that the participant reliably perceived. Empirically, we initialized *S*_0_ = 150 and Δ*S_t_* = 20.

### Near-threshold task

Following the one-up three-down procedure, participants performed several blocks of a near-threshold detection task (3 blocks for Participant 2, 4–6 blocks for Participants 3 and 4) in which the stimulus amplitude was fixed within each session. At this fixed intensity level, participants could sometimes perceive the stimulus and sometimes fail to detect it, yielding near-threshold variability in perception. No audible vibration sounds were present during the near-threshold task. If the selected amplitude consistently elicited either all “felt” or all “non-felt” responses, the threshold was re-calibrated using either a repeated one-up three-down staircase or manual adjustment to achieve a balanced distribution of perceptual outcomes. The default catch-trial proportion was set to 0.3 for Participants 2 and 3, who received higher-amplitude stimulation and exhibited low false alarm rates. For Participant 4, who had more intact sensation and a higher false alarm rate at lower input amplitudes, the catch-trial proportion was increased to 0.5. Of the 61 near-threshold sessions, 52 had valid accelerometer recordings and were included in force analyses.

### Ramping task

To examine the effect of physical stimulus amplitude on neural responses, a ramping task was conducted in separate sessions following the one-up three-down thresholding procedure. Because the highest ramping amplitudes produced audible mechanical sounds in some sessions, participants wore both noise-reducing earplugs and over-ear noise-isolating earmuffs during vibrotactile stimulation to eliminate auditory confounds associated with vibration. The task consisted of randomly presented trials across nine amplitude conditions: a catch condition with no stimulation (-100%), two sub-threshold levels (Sub1, 75% below the threshold amplitude; Sub2, -50%), one threshold level (0%), and five supra-threshold levels (Supra1–Supra5: +20%, +40%, +60%, +80% and +100%), with Supra1–Supra4 providing linearly spaced supra-threshold steps between threshold and Supra5. Supra5 corresponded to the highest amplitude that remained inaudible under the noise isolation setup. Sub1 and Sub2 were set to one-quarter and one-half of the threshold amplitude, respectively. On each trial, participants indicated whether they felt the stimulus and rated their perceived intensity on a scale from 1 to 10 after the fixation dot disappeared.

### Combined near-threshold and ramping task

To examine perceptual-state and amplitude coding in the same neurons, near-threshold and ramping tasks were combined within individual recording sessions in Participants 2, 3 and 4. Eight, five and six sessions were recorded in Participants 2, 3 and 4, respectively. Each session began with threshold estimation using the one-up three-down procedure, followed by 2–3 blocks each of the near-threshold and ramping tasks, yielding 80–120 near-threshold trials and 72–108 ramping trials per session. The order of the near-threshold and ramping tasks was counterbalanced across sessions. If the perceptual threshold changed during a session, the one-up three-down procedure was repeated to recalibrate the stimulation amplitude before subsequent blocks. To increase sample size, the near-threshold and ramping analyses also included data from the corresponding task blocks in these combined sessions.

### Vibrotactile discrimination task

To assess neural coding of physical stimulus amplitude while controlling variation in perceived intensity, we used a two-alternative forced-choice (2AFC) vibrotactile discrimination task as a control experiment. The tactor was placed on the index fingertip, where sensation was preserved and small force differences could be reliably distinguished. Before testing, we selected a fixed supra-threshold standard amplitude and identified lower and upper comparison amplitudes that could be confidently distinguished from the standard. The ranges between the standard and each endpoint were divided into five equal steps, yielding 11 comparison levels, including the standard. On each trial, cue onset was followed by a 1 s, 230 Hz standard stimulus, a 1 s delay and a 1 s comparison stimulus randomly selected from the 11 levels. After cue offset, the participant reported whether the comparison was stronger or weaker than the standard. The experiment was conducted in one participant (Participant 4) across three sessions. Each block comprised 44 trials, and each session included 6–8 blocks, yielding 264–352 trials per session.

### No-report control task

To dissociate percept-related neural activity from explicit reporting and associated motor responses, Participants 3 and 4 completed a near-threshold no-report control task. Following threshold estimation using the one-up three-down staircase, near-threshold blocks were organised as interleaved report and no-report blocks. Stimulation parameters, trial timing and catch-trial structure were identical across block types. In report blocks, participants indicated whether the vibration had been felt after the cue disappeared. In no-report blocks, no behavioural report was required, but participants were instructed to remain engaged and attend to whether the stimulus was perceived. Responses were compared across felt stimulus and non-felt stimulus trials in report blocks, stimulus trials in no-report blocks, and catch trials in both report and no-report blocks to determine whether percept-related modulation persisted without an overt report.

### Firing rate analysis

Firing rates were computed from spike-sorted data using 50 ms non-overlapping bins. To characterize the neural activity in response to the stimulus, the mean firing rate during the pre-stimulus baseline period (−2 to 0 s) was subtracted from the post-stimulus activity to yield baseline-corrected responses, i.e., the change in firing rate. For analyses based on firing rates across neurons, we used z-scored firing rates, in which each neuron was z-scored separately within each block by subtracting its mean firing rate and dividing by the corresponding standard deviation. For illustrative purposes, both single-neuron examples and population plots were smoothed using a causal Savitzky–Golay filter (polynomial order 1, frame length 11). Unless otherwise specified, all quantitative analyses were performed on the unsmoothed, baseline-corrected firing rates.

### Cell selection for the near-threshold task

To identify neurons encoding subjective sensory perception in the near-threshold task, we selected perception-sensitive neurons on a session-by-session basis. Neurons were defined as perception-sensitive if a one-way analysis of variance (ANOVA) revealed significant differences in baseline-corrected firing rates across stimulus-felt, stimulus-non-felt, and catch trial conditions. Two analysis windows were examined, i.e., the first 0.5 seconds and the second 0.5 seconds following stimulus onset. For Participant 4, due to a lower detection threshold and evidence accumulation of near-threshold detection, the analysis windows were shifted to 0–1 seconds and 1–2 seconds after stimulus onset. A neuron was considered perception-sensitive if there was a statistically significant difference (*P* < 0.05, FDR-corrected) in either of the two analysis windows.

### Cell selection for the ramping task

In the ramping task, neurons encoding stimulus amplitude were identified using a one-way ANOVA on firing rates across nine conditions, including five supra-threshold levels, one threshold level, two sub-threshold levels, and a catch trial condition. Two analysis windows were tested, spanning 0 to 0.5 seconds and 0.5 to 1 second after stimulus onset. A neuron was classified as sensory if there was a statistically significant difference (*P* < 0.05) in either window.

### Confidence cell selection

To examine how neural activity reflects subjective confidence, we analysed sessions from the near-threshold task in which participants provided a sufficiently varied distribution of confidence ratings. Sessions with uniform ratings were excluded. For felt trials, ratings were divided into high- and low-confidence groups using a session-specific threshold that ensured balanced trial counts. Since confidence ratings were not assessed in Participant 2, his data were excluded from this analysis. Neurons were considered confidence-sensitive if a one-way ANOVA revealed significant differences in baseline-corrected firing rates across high-confidence, low-confidence, and catch trial conditions. For non-felt trials, statistical significance was assessed using a one-way ANOVA to test for differences in baseline-corrected firing rates across confidence rating levels. Non-felt trials were similarly divided into high- and low-confidence groups using a session-specific threshold to balance trial counts. Due to insufficient non-felt trial counts, only data from Participant 4 were included in this analysis. Statistical tests were performed in two analysis windows (0–0.5 s and 0.5–1 s), and neurons were classified as confidence-sensitive if they reached significance (*P* < 0.05) in either window.

### Choice probability analysis

Choice probability quantifies a neuron’s ability to predict behavioural responses^53–55^, independent of model selection or parameter fitting used in machine learning based decoding approaches. It is derived from signal detection theory and reflects the separability of neural firing rate distributions across two behavioural outcomes, such as felt versus non-felt trials. Choice probability was quantified as the area under the curve (AUC) of the receiver operating characteristic (ROC), which compares the true positive and false positive rates across all possible decision thresholds. To reduce the confounding effects of noisy and task-unrelated neurons, choice probability was computed only for neurons previously identified as sensitive to perception or confidence. For perception-sensitive neurons, baseline-corrected firing rates from felt and non-felt trials were averaged over a 0–2 second window following stimulus onset to capture the full extent of subjective sensory perception preceding the report, and these values were used to compute choice probability. For confidence-sensitive neurons, choice probability was computed between high- and low-confidence trials.

For the ramping task, we applied the same ROC procedure to amplitude-sensitive neurons to compare firing rate distributions between amplitude conditions (e.g., Supra5 versus Supra1, Supra5 versus Supra2, or any stimulus level versus catch trials). We refer to this measure as AUC because it quantifies difference in stimulus amplitude rather than behavioural choice.

As both excitatory and inhibitory responses were observed across neurons, each neuron’s preferred response was used to define the positive class. Choice probability or AUC was computed for each neuron and pooled across sessions by brain region. To assess the temporal dynamics of encoding, we also computed time-resolved choice probability using a 200 ms sliding window. A null distribution was generated by randomly shuffling trial labels and recomputing choice probability or AUC 10 times, providing an empirical estimate of chance-level performance.

### Neuron-dropping analysis

To compare the informativeness of cortical areas in encoding perception or confidence using a population of single neurons, we performed a neuron-dropping analysis^56^ based on pseudo-populations. To aggregate data across sessions, firing rates were z-scored separately for each neuron before baseline correction. Neurons sensitive to the relevant variable were identified using these normalized responses and pooled across sessions within each cortical region. Their baseline-corrected firing rates during the stimulus period were used to construct the pseudo-population for decoding. To ensure fair comparisons across areas, we implemented a two-level subsampling procedure. First, we identified the minimum number of trials available across neurons and conditions (e.g., felt vs non-felt), and randomly sampled an equal number of trials per condition. Second, we incrementally reduced the number of neurons by randomly dropping one neuron at each step, repeating the classification using the remaining subset. At each step, a linear discriminant analysis (LDA) classifier with principal component analysis (PCA) for dimensionality reduction was trained on average firing rate changes in the 0–2 second window following stimulus onset to distinguish between the two conditions, and classification accuracy was obtained. Classification performance was evaluated using 10 repeated 70/30 train–test splits. PCA retained components explaining 80% of the variance, and the diagonal-linear LDA classifier was evaluated on the held-out trials. This process continued until only one neuron remained in the pseudo-population. Both the trial- and neuron-level subsampling loops were repeated 50 times, resulting in 2,500 subsampling iterations per area. Classification accuracy was averaged across cross-validation and subsampling repetitions. Unlike greedy selection methods that rank neurons by importance, our random sampling approach provided an unbiased estimate of the average information content per neuron in each cortical area. To enable fair comparisons across cortical areas, we identified the minimum number of available neurons and evaluated classification accuracy using matched neuron counts for each region.

### Single-neuron tuning analysis

To quantify the scale of cortical activation during near-threshold perception, we measured the proportion of neurons whose firing rates were modulated during felt or non-felt trials relative to correct-rejection catch trials. For each neuron and trial, firing rates in each response window were baseline-corrected using the pre-stimulus period. Neurons showing a significant difference between felt and catch trials or between non-felt and catch trials in either response window were classified as tuned to the corresponding condition. To account for unequal trial numbers, correct-rejection catch trials were randomly downsampled to match the number of felt and non-felt trials within each session, and this procedure was repeated 1,000 times; the resulting proportions of tuned neurons were averaged across resamples. For each cortical area, the proportion of tuned neurons was calculated within each session and then averaged across sessions.

### Analysis of the discrimination task

Neural responses were quantified as the mean denoised firing rate during the 1-s standard and comparison periods. Sensory-responsive neurons were selected based on stimulus-related modulation relative to the preceding 0.5-s baseline. Psychometric functions were fitted to trial-level behavioural choices as a function of comparison force using maximum-likelihood estimation. For each neuron, neural choices were defined by comparing responses to the comparison and standard stimuli, with response polarity reversed when firing decreased with force. Neurometric functions were fitted using the same procedure. Psychometric and neurometric just noticeable differences (JNDs) were calculated from the fitted 25% and 75% points, and neural-to-behavioural JND ratios were converted to decibels and grouped by cortical area.

For force decoding, trials were restricted to the psychometric JND range to focus on perceptually ambiguous stimuli for which both behavioural choices occurred. The central force levels were removed, and the remaining levels were divided into low- and high-force groups. Trials were resampled to match behavioural choice between force groups, and neuron counts were matched across areas. For choice decoding, trials in which the comparison was judged stronger or weaker than the standard were balanced within each force level, with neuron counts matched across areas. Pseudo-populations were constructed across sessions using 50 trial-resampling iterations and 50 matched-neuron resamplings per iteration. Linear discriminant analysis was evaluated using ten repeated cross-validation splits, and performance was quantified using held-out balanced accuracy.

### Neural latent dynamics

To examine how perception, amplitude, and confidence were encoded in the latent dynamics of neural activity, we visualized the temporal evolution of neural trajectories in a low-dimensional manifold. For each session, firing rates were z-scored per neuron and averaged across trials for neurons sensitive to the variable of interest. Smoothed firing rates were extracted from a window spanning −1 to 2 seconds relative to stimulus onset and pooled across sessions. To reduce noise and highlight neural patterns in the data, trial-averaged time series were projected onto the first three principal components using principal component analysis (PCA). This yielded low-dimensional trajectories that captured the dominant temporal dynamics for each condition within each cortical area.

To compare manifold structure across areas, we performed Procrustes alignment using activity from perception-sensitive neurons during felt trials in the near-threshold task. For each area, baseline-corrected felt, non-felt and catch responses were concatenated and projected onto a common three-dimensional PCA space, after which the felt trajectory was extracted for alignment. Each target-area trajectory was aligned to the corresponding S1, AIP or SMG reference trajectory at matched time bins. The Procrustes transformation estimated the rotation, scaling and translation that minimized the squared difference between the target and reference trajectories. Alignment quality was quantified using the normalized Procrustes disparity, and similarity was defined as one minus this value, with larger values indicating greater correspondence in trajectory geometry and temporal evolution. The aligned trajectories were also visualized in three-dimensional PCA space to compare their shapes and progression over time. Statistical significance was assessed using 1,000 permutations in which the temporal order of the target trajectory was randomly shuffled before realignment.

### Cross-session neuron overlap analysis

Because spike sorting was performed independently for each recording session, we assessed whether putatively identical neurons were recorded across sessions in Participants 3 and 4 using spike-waveform and inter-spike interval (ISI) similarity^57,58^. For each neuron, mean spike waveforms were mean-centered and normalized to unit length, and ISIs were restricted to finite positive values. Candidate cross-session pairs were limited to units recorded on the same electrode. Waveform similarity was quantified by waveform correlation and ISI similarity by the two-sample Kolmogorov–Smirnov (KS) statistic. Matching rate was defined as the proportion of valid same-electrode pairs classified as putative matches, and cross-session matching rates were visualized as a session-by-session heatmap. To determine whether repeated sampling influenced the main results, matched neurons were deduplicated by retaining their earliest occurrence and excluding matched occurrences from later sessions. Choice probability and population-decoding analyses were then repeated after removing these putatively overlapping neurons using the same neuronal selection and analysis parameters as in the original analyses.

### Cross-condition generalization analysis

To test whether perceptual representations generalized across the presence or absence of physical stimulation, we performed cross-condition decoding in Participant 4, who had sufficient false-alarm trials. Population decoding followed the same pseudo-population PCA–LDA procedure described in the neuron-dropping analysis, using all available neurons with matched trial and neuron counts across cortical areas. Two complementary cross-condition scenarios were tested: (1) the classifier was trained to distinguish hit from miss trials and tested on false alarm versus correct rejection trials, and (2) the classifier was trained on false alarm versus correct rejection trials and tested on hit versus miss trials. In both directions, the classifier was trained on one complete condition pair and evaluated on the other, providing an independent test set without an additional within-condition split. Trial resampling was repeated 50 times, with 50 matched-neuron subsamples per resampling, yielding 2,500 decoding estimates per cortical area. Classification accuracy was calculated for each repeat.

### Cross-time representational similarity analysis

To quantify the temporal stability of perceptual representations, we performed cross-time representational similarity analysis (RSA) using perception-sensitive activity from the near-threshold task. For each session and cortical area, area-specific pseudo-populations were constructed using balanced numbers of felt and non-felt trials. At each time bin, each trial was represented by a population activity vector containing the firing rates of the selected neurons. Pearson correlations were calculated between population activity vectors for every pair of time bins. The RSA contrast was defined as the mean correlation between trials with the same perceptual outcome minus the mean correlation between trials with different perceptual outcomes. Positive values therefore indicated that population activity patterns were more similar across time within the same perceptual state than across perceptual states. Session-level RSA contrast maps were averaged within each cortical area and participant to visualize the temporal evolution and persistence of perceptual representations.

### Modulation index for the combined near-threshold and ramping task

To quantify the relative encoding of perceptual choice and stimulus amplitude, we used a generalized linear model (GLM) incorporating both factors, *FR_stim_*∼*A_z_* + *P_stim_* + *C* + *FR_base_* + *t*, where *A_z_* and *P_stim_* denote standardized stimulus amplitude and perceptual report, respectively. *FR_stim_* and *FR_base_* denote firing rates during the stimulus and baseline periods, respectively. *C* and *t* denote the catch-trial indicator (0/1) and trial order, respectively. Perception tuning was identified using all near-threshold blocks, whereas amplitude tuning was evaluated using leave-one-block-out cross-validation within the ramping task. Neurons tuned to both factors were considered jointly tuned.

For each tuned neuron in either task, two reduced models were fitted to the ramping training blocks by separately excluding amplitude or perceptual report and used to predict firing rates in the held-out ramping block. Errors were averaged across folds, and the modulation index was defined as *MI* = *log*_10_(*SSE_noP_* / *SSE_noA_*), where *SSE_noP_* and *SSE_noA_* denote the prediction errors after excluding perceptual encoding and amplitude encoding, respectively. Positive MI values indicated stronger perceptual modulation, typically associated with nonlinear sigmoidal response profiles with saturation, whereas negative values indicated stronger amplitude modulation, typically associated with graded ramping responses across stimulus amplitudes.

### Single-trial activation and multi-area analyses

To quantify cortical recruitment on individual felt trials, single-trial neuronal activation was assessed using sorted z-scored firing rates. For each neuron, perception-related firing-rate changes on felt trials were compared with the distribution from non-felt trials using baseline (−2 to 0 s), stimulus-related response (0–2 s), and post-response (2–4 s) windows. A neuron was considered active when its response exceeded the non-felt mean by more than 2 standard deviations in either direction and subsequently returned toward baseline, defined as a post-response deviation within 1 standard deviation of the non-felt baseline and no greater than 50% of the stimulus-response deviation. Neurons required at least five valid non-felt trials for threshold estimation. An area was considered active when at least one neuron met these criteria. The activation probability was quantified as *P* (Activation Felt) for individual areas and all multi-area combinations. To test whether co-activation exceeded that expected from independent recruitment, the expected activation probability for each multi-area combination was calculated within each session as the product of the individual area activation probabilities, ∏*_i_ P_i_* (Activation ∣ Felt), and then averaged across sessions weighted by felt-trial count. For multi-area choice probability, perception-sensitive single-neuron responses were pooled across different cortical areas, and choice probability was calculated for each possible area combination.

### Neural latency analysis

To characterize neural latency across cortical areas, we analysed responses from the ramping and near-threshold tasks. For supra-threshold responses, latency was defined as the first time bin showing a significant firing-rate difference between the Supra5 and catch conditions. For near-threshold responses, neural divergence latency was defined as the first bin showing a significant difference between felt and non-felt trials, provided that significance persisted for at least three consecutive bins. Firing rates for the latency analysis were recomputed using 1-ms bins to improve temporal precision. Valid neuron-level latency estimates were pooled across sessions within each cortical area. Neurons without a finite latency estimate were excluded.

### Linear regression analysis

In the ramping task, we performed linear regression to assess the relationship between input force and neuronal firing rates. Input force was estimated from the recorded acceleration signals using the equation 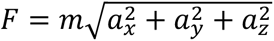, where *a_x_*, *a_y_*, and *a_z_*, represent acceleration along the three orthogonal axes, and *m* is the mass of the tactor. For each neuron, firing rates were regressed against force values across all amplitude levels, and both the slope and coefficient of determination (R²) were extracted to quantify the goodness-of-fit and the strength of the relationship.

To assess amplitude coding independent of conscious perception, a subliminal analysis was performed in Participant 2, whose higher vibrotactile detection threshold provided sufficient non-felt trials across stimulus amplitudes. Linear regression was repeated using only non-felt, non-catch trials. To assess amplitude coding while controlling for perceived intensity, a supraliminal analysis was performed in Participant 4, who had a lower detection threshold and sufficient felt trials across intensity levels. Force–firing rate relationships were evaluated within individual felt-intensity levels, thereby reducing covariation between physical amplitude and subjective intensity. For both analyses, R² and absolute slope values were compared across cortical areas.

### Statistics

Unless otherwise specified, comparisons between two conditions were performed using the non-parametric Mann–Whitney U (Wilcoxon rank-sum) test to avoid assumptions about underlying data distributions. Multiple comparisons were controlled using the Benjamini–Hochberg false discovery rate (FDR) procedure unless otherwise noted, with comparison families specified for each analysis in the Supplementary Information. Statistical significance is indicated by asterisks: *, *P* < 0.05; **, *P* < 0.01; and ***, *P* < 0.001.

For cases with limited session counts or sparse data, cluster-based non-parametric permutation testing was employed. When comparing more than two conditions, one-way ANOVA was used to assess the presence of significant differences across groups. To control for false positives in the identification of sensitive neurons, *P* values were corrected for multiple comparisons using FDR within each brain region. When too few neurons remained for population-level analysis, an uncorrected threshold of *P* < 0.05 was used.

To assess the significance of the number of neurons tuned to perception, amplitude, or confidence in each brain region, we constructed null distributions through permutation testing. Condition labels were randomly shuffled, and the full cell selection procedure was applied to compute the number of tuned neurons under the null hypothesis. This process was repeated 1,000 times. Statistical significance was determined by the proportion of permutations in which the number of tuned neurons exceeded the observed value.

To determine whether the observed proportion of tuned neurons exceeded chance levels, permutation-derived null distributions were generated using 10,000 permutations by shuffling trial-condition labels within each session and repeating the same tuning procedure. Empirical *P* values were calculated by comparing the observed area-level proportions with the corresponding null distributions, and multiple comparisons across cortical areas were controlled using FDR correction.

For cross-session neuron matching, null distributions of waveform and ISI similarity were generated from 5,000 randomly sampled neuron pairs recorded on different electrodes across sessions. Empirical *P* values were calculated separately for waveform and ISI similarity and combined using Fisher’s method. Same-electrode neuron pairs with a combined *P* < 0.01 were classified as putative cross-session matches.

For the modulation index analysis, perception-state and amplitude-modulated neurons were identified at FDR-corrected *P* < 0.05. Neurons classified as jointly tuned in at least 50% of ramping cross-validation folds were considered reliably jointly tuned.

For single- and multi-area activation analyses, statistical significance was assessed using 10,000 within-session permutations of felt, non-felt and catch labels, with the complete activation procedure recalculated after each permutation. Empirical *P* values were corrected for multiple comparisons using the FDR procedure. To test whether observed multi-area co-activation exceeded that expected under independent recruitment, trial order was independently shuffled for each area within each session 10,000 times, preserving marginal activation probabilities while disrupting trial-by-trial cross-area coordination. The observed co-activation rate was compared with this null distribution using a one-sided permutation test.

For the subliminal and supraliminal regression analyses, force values were shuffled 1,000 times to generate null distributions. R² and absolute slope values were compared with shuffled values using one-sided paired Wilcoxon signed-rank tests, and across cortical areas using two-sided Mann–Whitney U tests.

**Supplementary Table 1.** Recording sessions and neurons by task. Neuron counts represent spike-sorted units pooled across participants and recording sessions.

| Task | Participant | Sessions | Neurons |
|---|---|---|---|
| Near-threshold task | Participant 2 | 30 | 3,461 |
|  | Participant 3 | 15 | 5,703 |
|  | Participant 4 | 16 | 8,752 |
|  | **All** | **61** | **17,916** |
| Ramping task | Participant 2 | 6 | 653 |
|  | Participant 3 | 7 | 2,668 |
|  | Participant 4 | 5 | 2,447 |
|  | **All** | **18** | **5,768** |
| Combined near-threshold and ramping task | Participant 2 | 8 | 1,186 |
|  | Participant 3 | 5 | 2,544 |
|  | Participant 4 | 6 | 2,190 |
|  | **All** | **19** | **5,920** |
| Vibrotactile discrimination task | Participant 4 | 3 | 1,234 |
| No-report control task | Participant 3 | 1 | 469 |
|  | Participant 4 | 1 | 413 |
|  | **All** | **2** | **882** |
| **Total** | Participant 2 | 44 | 5,300 |
|  | Participant 3 | 28 | 11,384 |
|  | Participant 4 | 31 | 15,036 |
|  | **All** | **103** | **31,720** |

**Supplementary Table 2.** Post-hoc power of the comparisons between posterior parietal areas and S1. For each two-sided Wilcoxon rank-sum test (α = 0.05) comparing task-modulated neurons of a posterior parietal area with those of S1, the table reports the recorded numbers of neurons (area / S1), the minimal numbers required to achieve 80% power while maintaining the recorded ratio of area to S1 neurons, and the estimated statistical power at the recorded numbers of neurons. Power estimates used the observed effect sizes (rank-biserial correlation |*r*|, Supplementary Table 3). Near-threshold task: choice probability of single neurons for felt versus non-felt trials (Fig. 2e–g). Ramping task: R² of the linear regression of each tuned neuron’s response on stimulation force (Fig. 3i). Estimated power exceeded 80% for all comparisons (82.5% to >99.9%), indicating sufficient power at the neuronal level to detect the observed differences.

| Participant | Task | Comparison | Recorded neurons | Estimated minimum neurons for 80% power (area / S1) | Statistical power (%) |
|---|---|---|---|---|---|
| **Participant 2** | Near-threshold | SMG vs S1 | 36 / 234 | 19 / 120 | 97.6 |
|  | Ramping | SMG vs S1 | 24 / 135 | 8 / 42 | 99.9 |
| **Participant 3** | Near-threshold | AIP vs S1 | 368 / 181 | 345 / 170 | 82.5 |
|  | Near-threshold | SMG vs S1 | 413 / 181 | 157 / 69 | 99.5 |
| **Participant 4** | Near-threshold | AIP vs S1 | 269 / 228 | 164 / 139 | 94.9 |
|  | Near-threshold | SMG vs S1 | 446 / 228 | 46 / 24 | >99.9 |
|  | Ramping | AIP vs S1 | 193 / 199 | 144 / 149 | 90.0 |
|  | Ramping | SMG vs S1 | 104 / 199 | 14 / 26 | >99.9 |

**Supplementary Table 3.** Sensitivity and effect sizes of the comparisons between posterior parietal areas and S1. For each two-sided Wilcoxon rank-sum test (α = 0.05), the table reports the smallest rank-biserial correlation |r| detectable with 80% power at the recorded numbers of neurons (Supplementary Table 2), the observed effect size |r| = |2A − 1|, and its bootstrap 95% confidence interval (10,000 resamples of the neurons), where A is the probability that a neuron of the posterior parietal area has a higher value than a neuron of S1. Choice probability was higher in the posterior parietal areas than in S1 (near-threshold task), and R² was lower (ramping task). All observed effect sizes exceeded the corresponding detection thresholds for 80% power, indicating sufficient sensitivity at the neuronal level for these comparisons.

| **Participant** | **Task** | **Comparison** | **Smallest detectable \|*r*\| (80% power)** | **\|*r*\|** | **95% CI** |
|---|---|---|---|---|---|
| **Participant 2** | Near-threshold | SMG vs S1 | 0.290 | 0.408 | [0.216, 0.588] |
|  | Ramping | SMG vs S1 | 0.359 | 0.644 | [0.478, 0.790] |
| **Participant 3** | Near-threshold | AIP vs S1 | 0.147 | 0.152 | [0.051, 0.251] |
|  | Near-threshold | SMG vs S1 | 0.144 | 0.234 | [0.135, 0.328] |
| **Participant 4** | Near-threshold | AIP vs S1 | 0.146 | 0.187 | [0.086, 0.284] |
|  | Near-threshold | SMG vs S1 | 0.132 | 0.413 | [0.332, 0.491] |
|  | Ramping | AIP vs S1 | 0.164 | 0.189 | [0.075, 0.302] |
|  | Ramping | SMG vs S1 | 0.196 | 0.549 | [0.443, 0.650] |

## Additional details for figures

Figure 1a. Data from the ventral premotor cortex (PMv) array in Participant 2 were not analysed because of substantial degradation in single-neuron signals.

Figure 1c. A vibrotactile armband delivered stimulation to sensate regions (bicep, finger, or elbow), while intracortical single-neuron activity was simultaneously recorded via Utah arrays connected to three percutaneous pedestals.

Figure 1d. Each trial began with a fixation dot, followed by a variable delay (2–3 s) before a 1 s vibrotactile stimulus (230 Hz). After a 1.5 s delay, the fixation dot disappeared, prompting participants to report whether they perceived the stimulus (felt/non-felt). Participants 3 and 4 additionally rated their confidence (1–10). Catch trials (30–50%) without stimulation were included to control for false detections.

Figure 1e. The schematic illustrates how stimulus amplitude increased after a non-felt report and decreased after three consecutive felt reports, adaptively converging on the vibrotactile detection threshold.

Figure 1g. At near-threshold force levels, stimuli can be reported as either felt or non-felt despite comparable physical input.

Figure 1h. Nine vibrotactile amplitudes, including five supra-threshold levels (+20%, +40%, +60%, +80%, and +100%), one threshold (0%), two sub-threshold conditions (- 75%, -50%), and a catch trial with no stimulation, were randomly presented during the ramping task. The sub-threshold levels were set to 25% and 50% of the threshold amplitude. Positive percentage labels denote linearly spaced amplitude settings between threshold (0%) and the maximum inaudible amplitude (+100%), rather than percentage increases in measured force.

Figure 1i. Heatmaps contained *n* = 333/356/451/307 neurons in S1/AIP/SMG/M1, respectively, pooled across participants. For each participant, neuron counts were *n* = 146/20 in S1/SMG for Participant 2, 87/231/191/123 in S1/AIP/SMG/M1 for Participant 3, and 100/125/240/184 in S1/AIP/SMG/M1 for Participant 4. Neurons were selected for significant positive sensory modulation after Benjamini–Hochberg FDR correction.

Figure 1j. Felt trials recruited sensitive neurons across multiple cortical areas, whereas significant tuning during non-felt trials was observed in S1 and to a smaller extent in AIP in one participant. For felt trials, Participant 2: S1, *P* = 3.0 × 10^−^⁴; SMG, *P* = 0.0050. Participant 3: AIP, SMG and M1, all *P* = 3.3 × 10^−^⁴. Participant 4: S1, AIP, SMG and M1, all *P* = 2.5 × 10^−^⁴; dlPFC, *P* = 0.0086. For non-felt trials, mean tuned-neuron percentages across sessions were 3.51% in Participant 2’s S1, *P* = 3.0 × 10^−^⁴, and 0.76% in Participant 3’s AIP, *P* = 0.0042; all *P* values are Benjamini–Hochberg FDR-adjusted. One-sided permutation tests against a catch-matched null used 10,000 permutations.

Figure 2a. Traces show normalized firing rates averaged across perception-sensitive neurons for the three participants. Perception-sensitive cells were identified by the firing rates using one-way ANOVA across felt, non-felt, and catch conditions.

Figure 2b. Firing rates of PPC and S1 neurons increased in the felt condition both when vibration was delivered and during catch trials without stimulation (error trials / false alarm).

Figure 2c. As no stimulus was delivered, tactile perception could occur at variable times, and neural signals were not time-locked to perception onset. Data from Participant 4 are shown due to a higher rate of false alarms associated with a lower vibrotactile threshold. Figure 2d. Choice probability is calculated per neuron as the area under the receiver operating characteristic (ROC) curve (bottom) and reflects the separation between firing rate distributions for the two perceptual conditions (top).

Figure 2e–g. Sample sizes for choice probability: *n* = 36/234 cells in SMG/S1 for Participant 2, and 368/413/181 and 269/446/228 cells in AIP/SMG/S1 for Participants 3 and 4, respectively. Sample sizes for neuron-dropping insets: *n* = 34/181/169 cells per area for Participants 2/3/4, respectively. Each dot denotes the mean decoding accuracy from one of 50 trial-resampling repetitions, averaged across 50 neuron subsamples and 10 train–test splits per subsample. Bars show means and error bars indicate 95% confidence intervals across repetitions. Neurons for neuron-dropping analysis were selected separately using z-scored firing rates (Methods). For Participants 3 and 4, Benjamini–Hochberg FDR correction was applied across the two PPC–S1 comparisons.

Figure 2h. Sample sizes for choice probability: *n* = 107/172/73 cells in AIP/SMG/S1; Neuron-dropping insets: *n* = 178 cells. *P* values are Benjamini–Hochberg FDR-adjusted across the two PPC–S1 comparisons.

Figure 2i. Crosses mark trial onset; red and blue indicate stimulus onset, and yellow and green indicate stimulus offset. Neural trajectories in latent space diverge between felt and non-felt conditions, indicating that subjective awareness is encoded in lower-dimensional activity patterns beyond average firing rates. Neural trajectories included *n* = 35/209 perception-sensitive neurons in SMG/S1 for Participant 2, and 207/91/12 and 48/202/22 neurons in AIP/SMG/S1 for Participants 3 and 4, respectively.

Figure 3a. Heatmaps contained n = 410/453/170 positively responding neurons in S1/AIP/SMG, respectively, pooled across Participants 2–4. Positive percentage labels denote stimulus-amplitude settings scaled between threshold (0%) and the maximum inaudible amplitude (+100%). Stronger vibrotactile amplitudes evoked increased neural activation in PPC and S1.

Figure 3b. Firing rate increased with applied force, and significant correlations were observed, particularly in S1. Each of the three example neurons included *n* = 216 trials, with 24 trials per force condition. Force–firing rate correlations were significant for all three neurons (all *P* < 10^−^²⁴; two-tailed Pearson tests).

Figure 3c. Normalized firing rates across the nine input amplitudes are shown for PPC and S1 in each participant. Firing rates increased with stimulus amplitude, with a more prominent increase in S1 for Participant 2 and Participant 4. This analysis and all subsequent analyses used task-modulated neurons, identified via ANOVA based on significant differences in firing across the nine amplitude conditions. Neurons with significantly increased firing at the highest amplitude relative to baseline were included in this panel. X-axis labels denote catch trials (C), the two sub-threshold levels at 25% and 50% of the threshold amplitude (S1 and S2, respectively), non-felt and felt trials at threshold (T− and T+, respectively), and five supra-threshold levels scaled between threshold and the maximum inaudible amplitude (S1+ to S5+, corresponding to +20%, +40%, +60%, +80% and +100%, respectively). On the x-axis, S1 and S2 denote Sub1 and Sub2, respectively, and S1+–S5+ denote Supra1–Supra5. Neuronal sample sizes: *n* = 29/191 neurons in SMG/S1 for Participant 2, and 245/102/54 and 237/123/261 neurons in AIP/SMG/S1 for Participants 3 and 4, respectively.

Figure 3d. AUC was computed by comparing firing rates at each amplitude level to those during catch trials. A general increase in AUC with stimulus amplitude was observed. T denotes threshold stimulation; the remaining x-axis abbreviations are defined in Fig. 3c. Neuronal sample sizes: *n* = 74/191 neurons in SMG/S1 for Participant 2, and 306/301/138 and 319/227/434 neurons in AIP/SMG/S1 for Participants 3 and 4, respectively. Felt and non-felt trials were not split at each amplitude to avoid increased noise in AUC estimates from smaller trial counts.

Figure 3e,f. S1 exhibited higher AUC values than PPC for distinguishing Supra5 from Supra1, indicating greater sensitivity to amplitude differences. Participant 2: *P*_SMG,S1_ = 4.6 × 10^−^⁷. Participant 4: *P*_AIP,S1_ = 2.9 × 10^−^⁴; *P*_SMG,S1_ = 2.5 × 10^−^¹¹. AUC distributions were compared using two-tailed Wilcoxon rank-sum tests. Participant 4’s reported P values are Benjamini–Hochberg FDR-adjusted across the two PPC–S1 comparisons. Neuronal sample sizes: *n* = 79/201 neurons in SMG/S1 for Participant 2 and 255/194/321 neurons in AIP/SMG/S1 for Participant 4.

Figure 3g,h. AUC was computed between +100% and +40% amplitudes for Participant 2 and Participant 4. Left panels show the distribution of AUC for PPC and S1 neurons, and right panels show the temporal profile computed in sliding bins. S1 exhibited higher AUC values than PPC for distinguishing Supra5 from Supra2, indicating greater sensitivity to amplitude differences. Participant 2: *P*_SMG,S1_ = 5.0 × 10^−^⁵. Participant 4: *P*_AIP,S1_ = 0.0155; *P*_SMG,S1_ = 2.4 × 10^−^⁸. AUC distributions were compared using two-tailed Wilcoxon rank-sum tests. Participant 4’s reported *P* values are Benjamini–Hochberg FDR-adjusted across the two PPC–S1 comparisons. Although the difference in AUC was smaller than that observed for +100% vs +20%, S1 consistently showed higher AUC values than PPC, indicating greater sensitivity to amplitude changes. Sample sizes are the same as Figure 3e,f.

Figure 3i. R^2^ values from linear regressions between firing rate and input force are shown for Participant 2 and Participant 4, with each dot representing a single neuron. Participant 4’s reported *P* values are Benjamini–Hochberg FDR-adjusted across the two PPC–S1 comparisons. Neuronal sample sizes: n = 24/135 neurons in SMG/S1 for Participant 2 and 193/104/199 neurons in AIP/SMG/S1 for Participant 4. The result indicates stronger linear relationships between firing rate and stimulus amplitude.

Figure 3j. Slope values from linear regressions between firing rate and input force are shown for Participant 2 and Participant 4, with each dot representing a single neuron. Sample sizes are the same as Figure 3i. Participant 4’s reported *P* values are Benjamini– Hochberg FDR-adjusted across the two PPC–S1 comparisons.

Figure 3k. For Participant 4, population trajectories included n = 237/123/261 amplitude-sensitive neurons in AIP/SMG/S1, respectively. Neural trajectories corresponding to different vibrotactile amplitudes show a clear gradient, with higher input amplitudes producing greater divergence in activity patterns during the ramping task.

Figure 3l. Latency was defined as the first time bin showing a significant difference in firing rate between the Supra5 and catch conditions. Comparisons were performed using two-tailed Wilcoxon rank-sum tests. Mean latency across participants showed that S1 responded earlier than PPC during supra-threshold stimulation. Sample sizes for Participant 2, *n* = 66/194 cells in SMG/S1; Sample sizes for Participant 3, *n* = 258/237/72 cells in AIP/SMG/S1. Sample sizes for Participant 4, *n* = 269/174/344 cells in AIP/SMG/S1. Participant 2: *P*_SMG,S1_ = 7.0 × 10^−^²¹. Participant 3: *P*_AIP,S1_ = 0.017; *P*_SMG,S1_ = 0.0017. Participant 4: *P*_AIP,S1_ = 0.010; *P*_SMG,S1_ = 6.1 × 10^−^²⁶. PPC–S1 *P* values for Participants 3 and 4 are Benjamini–Hochberg FDR-adjusted across the two comparisons within each participant.

Figure 4a. The example firing rate panel shows higher confidence was associated with increased firing rates. Participant 2 was not used for the confidence analyses because confidence was not assessed in this participant.

Figure 4b. Normalized firing rates are shown for Participant 3 and Participant 4, comparing high-confidence, low-confidence, and catch conditions. Confidence-tuned neurons were identified via ANOVA across confidence levels in the near-threshold task. Neuronal sample sizes: *n* = 109/87/47 and 76/98/97 neurons in AIP/SMG/S1 for Participants 3 and 4, respectively.

Figure 4c,d. Choice probability was computed for confidence-tuned neurons in Participant 3 and Participant 4. Sample sizes for Participant 3, *n* = 80/141/67 cells in AIP/SMG/S1; Sample sizes for Participant 4, *n* = 73/116/91 cells in AIP/SMG/S1. *P* values are Benjamini–Hochberg FDR-adjusted across the two PPC–S1 comparisons.

Figure 4e. Choice probability was computed for confidence-tuned neurons in Participant 4, comparing high- and low-confidence non-felt trials. Participant 3 was not included due to an insufficient number of balanced non-felt trials across confidence levels. Sample sizes for Participant 4, *n* = 106/101/152 cells in AIP/SMG/S1. *P* values are Benjamini– Hochberg FDR-adjusted across the two PPC–S1 comparisons.

Figure 4f. An LDA classifier was trained to distinguish high- and low-confidence felt trials using increasing numbers of neurons. The result of neuron-dropping analysis is consistent with single-neuron choice probability results in panels c and d. The inset displays classification accuracy at the neuron count marked by the dashed line. Insets used 128/96 neurons per area for Participants 3/4, respectively, with 50 decoder repetitions per area.

Figure 4g. An LDA classifier was trained to distinguish high- and low-confidence non-felt trials using increasing numbers of neurons. The result is consistent with the single-neuron choice probability result in panel e. Sample sizes for Participant 4, *n* = 89/97/124 cells in AIP/SMG/S1. The inset used 89 neurons per area, with 50 decoder repetitions per area. Figure 4h. PPC exhibited a higher proportion of mixed-selective neurons compared with S1, indicating stronger joint encoding of perception and confidence.

Figure 4i. Neural trajectories from Participant 3 and Participant 4 show distinct neural dynamics for high- and low-confidence felt trials, reflecting confidence encoding in the neural manifold. Warmer colours represent high-confidence felt trials, cooler colours represent low-confidence felt trials, and black represents catch trials. Crosses indicate trial onset; red and blue mark stimulus onset, and yellow and green mark stimulus offset.

Extended Data Figure 1g. Thin grey lines connect the mean forces measured by accelerometer for felt and non-felt trials within individual recording sessions. The thick grey line and large circles indicate the means across sessions. Participant 2: *P* = 0.89, *n* = 30 sessions; Participant 3: *P* = 0.84, *n* = 11 sessions; Participant 4: *P* = 0.65, *n* = 11 sessions; two-tailed rank-sum test on session means.

Extended Data Figure 1h. Participant 3: *P* = 1.00, *n* = 6 sessions; Participant 4: *P* = 0.38, n = 5 sessions; two-tailed rank-sum test on session means. In g and h, session counts denote sessions with valid accelerometer recordings.

Extended Data Figure 1i. Neuronal sample sizes: *n* = 410/453/170/186 cells in S1/AIP/SMG/M1.

Extended Data Figure 2a,b. Sample sizes for neurons with excitation responses in the ramping task: Participant 2, *n* = 175/10 cells in S1/SMG; Participant 3, *n* = 20/277/88/32/5 cells in S1/AIP/SMG/M1/PMv; Participant 4, *n* = 215/176/72/154/3 cells in S1/AIP/SMG/M1/dlPFC. The neurons with excitation and inhibition responses refer to increases and decreases in firing rate, respectively, and do not indicate excitatory or inhibitory neuronal cell types.

Extended Data Figure 2c. Neuronal sample sizes: Participant 2, *n* = 213/35 cells in S1/SMG; Participant 3, *n* = 25/317/140/72/6 cells in S1/AIP/SMG/M1/PMv; Participant 4, *n* = 303/231/96/210/8 cells in S1/AIP/SMG/M1/dlPFC. Combined-task sessions were also included. Across all cortical regions and participants, no shuffled count exceeded the observed number of tactile-tuned neurons, as assessed using a one-sided permutation test.

Extended Data Figure 2h. Sample sizes for neurons with inhibition responses in the ramping task: Participant 2, *n* = 46/31 cells in S1/SMG; Participant 3, *n* = 5/40/52/40 cells in S1/AIP/SMG/M1; Participant 4, *n* = 88/55/24/56 cells in S1/AIP/SMG/M1.

Extended Data Figure 3a. Sample sizes for neurons with excitation responses in the near-threshold task: Participant 2, *n* = 146/20 cells in S1/SMG; Participant 3, *n* = 87/231/191/123 cells in S1/AIP/SMG/M1; Participant 4, *n* = 100/125/240/184/15 cells in S1/AIP/SMG/M1/dlPFC. Also, excitation and inhibition describe increases and decreases in firing rate, respectively, not neuronal cell types.

Extended Data Figure 3b. Neuronal sample sizes: Participant 2, *n* = 225/37 cells in S1/SMG; Participant 3, *n* = 181/368/413/252/49 cells in S1/AIP/SMG/M1/PMv; Participant 4, *n* = 228/269/446/439/20 cells in S1/AIP/SMG/M1/dlPFC. For Participant 3, *P* = 0.168 for PMv, using one-sided permutation tests. For all other displayed regions and participants, no shuffled count exceeded the observed count across 1,000 shuffles.

Extended Data Figure 3c. Neuronal sample sizes: Participant 3, *n* = 181/368/413/252/49 cells in S1/AIP/SMG/M1/PMv. Before correction, Participant 3: *P*_SMG,S1_ = 5.4 × 10^−6^, *P*_AIP,S1_ = 0.0038, *P*_M1,S1_ = 0.496, *P*_PMv,S1_ = 0.788; two-tailed rank-sum tests. After Benjamini–Hochberg FDR correction across four comparisons, Participant 3: *P*_SMG,S1_ = 2.1 × 10^−5^, *P*_AIP,S1_ = 0.0076, *P*_M1,S1_ = 0.661, *P*_PMv,S1_ = 0.788.

Extended Data Figure 3d. Neuronal sample sizes: Participant 4, *n* = 222/266/430/430/17 cells in S1/AIP/SMG/M1/dlPFC. Before correction, Participant 4: *P*_SMG,S1_ = 5.3 × 10^−19^, *P*_AIP,S1_ = 1.57 × 10^−4^, *P*_M1,S1_ = 0.00107, *P*_dlPFC,S1_ = 0.0027; two-tailed rank-sum tests. After Benjamini–Hochberg FDR correction across four comparisons, Participant 4: *P*_SMG,S1_ = 2.1 × 10^−18^, *P*_AIP,S1_ = 3.1 × 10^−4^, *P*_M1,S1_ = 0.0014, *P*_dlPFC,S1_ = 0.0027.

Extended Data Figure 3e. Pseudo-populations were matched at 49 neurons per area for Participant 3. Before correction, *P*_SMG,S1_ = 5.0 × 10⁻¹⁶, *P*_AIP,S1_ = 1.2 × 10⁻¹⁰, *P*_M1,S1_ = 0.00024; two-tailed rank-sum tests on decoding accuracy at the matched neuron count. After Benjamini–Hochberg FDR correction across four comparisons, *P*_SMG,S1_ = 2.0 × 10⁻¹⁵, *P*_AIP,S1_ = 2.4 × 10⁻¹⁰, *P*_M1,S1_ = 0.00032, *P*_PMv,S1_ = 0.011.

Extended Data Figure 3f. Pseudo-populations were matched at 14 neurons per area. Before correction, *P*_SMG,S1_ = 7.1 × 10^−^¹⁸, *P*_AIP,S1_ = 7.1 × 10^−^¹⁸, *P*_M1,S1_ = 2.0 × 10^−^¹⁷, *P*_dlPFC,S1_ = 1.4 × 10^−^¹⁴; two-tailed rank-sum tests on decoding accuracy at the matched neuron count. After Benjamini–Hochberg FDR correction across four comparisons, *P*_SMG,S1_ = 1.4 × 10^−^¹⁷, *P*_AIP,S1_ = 1.4 × 10^−^¹⁷, *P*_M1,S1_ = 2.6 × 10^−^¹⁷, *P*_dlPFC,S1_ = 1.4 × 10^−^¹⁴.

Extended Data Figure 3g. For false-alarm (FA) versus correct-rejection (CR) decoding, pseudo-populations were matched at 103 neurons per area. Before correction, *P*_SMG,S1_ = 9.1 × 10^−^⁴, *P*_AIP,S1_ = 6.7 × 10^−^¹⁵, *P*_M1,S1_ = 5.3 × 10^−^⁷, *P*_dlPFC,S1_ = 1.5 × 10^−^⁴; two-tailed rank-sum tests on decoding accuracy. After Benjamini–Hochberg FDR correction across four comparisons, *P*_SMG,S1_ = 9.1 × 10^−^⁴, *P*_AIP,S1_ = 2.7 × 10^−^¹⁴, *P*_M1,S1_ = 1.1 × 10^−^⁶, *P*_dlPFC,S1_ = 2.1 × 10^−^⁴.

Extended Data Figure 3h. For cross-condition decoding, classifiers were trained on hit versus miss trials and tested on false-alarm (FA) versus correct-rejection (CR) trials. Pseudo-populations were matched at 103 neurons per area. Before correction, Participant 4: *P*_SMG,S1_ = 1.7 × 10^−^⁹, *P*_AIP,S1_ = 3.3 × 10^−^¹⁵, *P*_M1,S1_ = 5.6 × 10^−^¹², *P*_dlPFC,S1_ = 9.8 × 10^−^⁹; two-tailed rank-sum tests on decoding accuracy. After Benjamini–Hochberg FDR correction across four comparisons, *P*_SMG,S1_ = 2.2 × 10^−^⁹, *P*_AIP,S1_ = 1.3 × 10^−^¹⁴, *P*_M1,S1_ = 1.1 × 10^−^¹¹, *P*_dlPFC,S1_ = 9.8 × 10^−^⁹.

Extended Data Figure 3i. For cross-condition decoding, classifiers were trained on false-alarm (FA) versus correct-rejection (CR) trials and tested on hit versus miss trials. Pseudo-populations were matched at 103 neurons per area. Before correction, *P*_SMG,S1_ = 3.0 × 10^−^¹⁰, *P*_AIP,S1_ = 5.8 × 10^−^¹⁶, *P*_M1,S1_ = 1.4 × 10^−^¹³, *P*_dlPFC,S1_ = 1.7 × 10^−^⁶; two-tailed rank-sum tests on decoding accuracy. After Benjamini–Hochberg FDR correction across four comparisons, *P*_SMG,S1_ = 4.0 × 10^−^¹⁰, *P*_AIP,S1_ = 2.3 × 10^−^¹⁵, *P*_M1,S1_ = 2.8 × 10^−^¹³, *P*_dlPFC,S1_ = 1.7 × 10^−^⁶.

Extended Data Figure 3j. For choice-probability analysis after cross-session deduplication in j,k, the earliest occurrence of each putatively overlapping neuron was retained. Neuronal sample sizes for Participant 3, *n* = 148/227/208/170/28 cells in S1/AIP/SMG/M1/PMv, pooled across sessions. Before correction, *P*_SMG,S1_ = 7.1 × 10⁻⁵, *P*_AIP,S1_ = 0.0016, *P*_M1,S1_ = 0.51, *P*_PMv,S1_ = 0.19; two-tailed rank-sum tests on single-neuron choice probability. After Benjamini–Hochberg FDR correction across four comparisons, *P*_SMG,S1_ = 0.00028, *P*_AIP,S1_ = 0.0033, *P*_M1,S1_ = 0.51, *P*_PMv,S1_ = 0.26.

Extended Data Figure 3k. For choice-probability analysis after cross-session deduplication, the earliest occurrence of each putatively overlapping neuron was retained. Neuronal sample sizes for Participant 4, *n* = 152/146/193/280/13 cells in S1/AIP/SMG/M1/dlPFC, pooled across sessions. Before correction, Participant 4: *P*_SMG,S1_ = 4.6 × 10^−^¹¹, *P*_AIP,S1_ = 0.023, *P*_M1,S1_ = 0.0068, *P*_dlPFC,S1_ = 0.061; two-tailed rank-sum tests on single-neuron choice probability. After Benjamini–Hochberg FDR correction across four comparisons, Participant 4: *P*_SMG,S1_ = 1.8 × 10^−^¹⁰, *P*_AIP,S1_ = 0.031, *P*_M1,S1_ = 0.014, *P*_dlPFC,S1_ = 0.061.

Extended Data Figure 4c. Thin grey lines connect the mean forces measured by accelerometer for felt and non-felt trials within individual recording sessions, shown separately for each stimulation site in Participant 2. The thick grey line and large circles indicate the means across sessions. Elbow: *P* = 0.84, *n* = 14 sessions; bicep: *P* = 1.00, n = 7 sessions; finger: *P* = 0.90, *n* = 7 sessions; two-tailed rank-sum test on session means. Extended Data Figure 4d. For Participant 4, finger: *P* = 0.55, *n* = 5 sessions; bicep: *P* = 0.82, *n* = 6 sessions; two-tailed rank-sum test on session means. In c and d, session counts denote sessions with valid accelerometer recordings.

Extended Data Figure 4e. Bars show mean ± s.e.m. across sessions, and each dot denotes one session. For Participant 2, elbow, finger and biceps stimulation included 19, 7 and 10 sessions, respectively, including combined-task sessions. Recorded-neuron sample sizes were n = 1002/914 cells in S1/SMG for elbow stimulation, 544/307 for finger stimulation and 725/481 for biceps stimulation. Corresponding modulated-neuron counts were 237/90, 95/28 and 108/42, respectively. Participant 2: *P*_SMG_ = 0.82, *P*_S1_ = 0.036; Kruskal–Wallis tests comparing session-wise proportions across the three stimulation sites. For Participant 4, recorded-neuron sample sizes were *n* = 1173/666/651 and 1139/646/620 cells in S1/AIP/SMG for finger and biceps stimulation, respectively. Corresponding modulated-neuron counts were 273/158/204 and 165/134/124, respectively. Permutation tests were used for Participant 4 given the small number of sessions and the comparison of only two stimulation sites. Participant 4: *P*_AIP_ = 0.50, *P*_SMG_ = 0.066, *P*_S1_ < 5 × 10^−^⁵; paired permutation tests comparing session-wise proportions between the two stimulation sites, using 20,000 permutations. Counts were pooled across sessions within each stimulation site.

Extended Data Figure 4f. Single-neuron choice probabilities across stimulation sites. Each dot denotes one near-threshold modulated neuron with a choice probability. Boxes show the interquartile range, horizontal lines indicate medians, and whiskers extend up to 1.5 times the interquartile range. For Participant 2, neuronal sample sizes were *n* = 150/24 cells in S1/SMG for elbow stimulation, 45/5 for finger stimulation and 39/7 for biceps stimulation. Participant 2: *P*_SMG_ = 0.23, *P*_S1_ = 0.045; Kruskal–Wallis tests comparing single-neuron choice probabilities across the three stimulation sites. For Participant 4, neuronal sample sizes were *n* = 137/122/169 and 38/67/172 cells in S1/AIP/SMG for finger and biceps stimulation, respectively. Participant 4: *P*_AIP_ = 0.42, *P*_SMG_ = 0.30, *P*_S1_ = 0.0078; two-tailed rank-sum tests comparing single-neuron choice probabilities between the two stimulation sites. Neuron counts represent spike-sorted units pooled across sessions within each stimulation site.

Extended Data Figure 4g. Neuron-dropping decoding across stimulation sites in Participant 2. The denotation is the same as Figure 2e–g. Neuronal sample sizes were *n* = 10/102 cells in SMG/S1 for elbow stimulation, 4/39 for finger stimulation and 7/38 for biceps stimulation. Insets show decoding accuracy with neuron numbers matched across areas, as indicated by the dashed lines: 10, 4 and 7 neurons per area for elbow, finger and biceps stimulation, respectively. SMG versus S1: elbow, *P* = 1.1 × 10^−^⁹; biceps, *P* = 1.9 × 10^−^¹²; two-tailed rank-sum tests. For finger stimulation, only four SMG neurons were available; the comparison at four neurons per area used a two-tailed paired permutation test with 20,000 permutations, *P* = 0.044.

Extended Data Figure 4h. Neuron-dropping decoding across stimulation sites in Participant 4. Neuronal sample sizes were *n* = 93/127/122 and 78/74/32 cells in AIP/SMG/S1 for finger and biceps stimulation, respectively. Matched neuron counts in insets: 93 and 32 neurons per area for finger and biceps stimulation, respectively. Finger stimulation: *P*_AIP,S1_ = 7.1 × 10^−^¹⁸, *P*_SMG,S1_ = 7.1 × 10^−^¹⁸; biceps stimulation: *P*_AIP,S1_ = 1.8 × 10^−^¹¹, *P*_SMG,S1_ = 7.5 × 10^−^¹⁷; two-tailed rank-sum tests comparing decoding-repeat accuracies. Neuron counts represent spike-sorted units pooled across sessions within each stimulation site in g,h.

Extended Data Figure 6a. For the combined near-threshold and ramping task, sample sizes with/without near-threshold modulation were n = 16/331 cells in SMG and 51/256 in S1 for Participant 2, and 26/387 in AIP, 30/400 in SMG, 29/528 in S1 and 47/442 in M1 for Participant 4. Participant 2: *P*_SMG_ = 0.013, *P*_S1_ = 2.2 × 10^−^¹¹; Participant 4: *P*_AIP_ = 0.0016, *P*_SMG_ = 0.013, *P*_S1_ = 0.0014, *P*_M1_ = 0.0015; two-tailed rank-sum tests comparing neurons with versus without near-threshold modulation within each region. Each dot denotes one neuron; open circles and error bars show mean ± s.d.

Extended Data Figure 6b. Amplitude-modulated / total neuron counts in groups with versus without near-threshold modulation were: Participant 2, SMG, 11/16 versus 119/331; S1, 41/51 versus 103/256. Participant 4, AIP, 12/26 versus 80/389; SMG, 11/30 versus 57/407; S1, 19/30 versus 144/594; M1, 25/47 versus 135/443.

Extended Data Figure 6d. For Participant 2, the modulation index comparison included *n* = 141/162 cells in SMG/S1. *P*_SMG,S1_ = 6.5 × 10^−^⁵; two-tailed rank-sum test.

Extended Data Figure 6e. For Participant 4, neuronal sample sizes in the modulation index comparison panel were *n* = 106/87/174 cells in AIP/SMG/S1. *P*_AIP,S1_ = 0.49, *P*_SMG,S1_ = 0.0012; two-tailed rank-sum tests.

Extended Data Figure 6f. For Participant 3, who received stimulation at a site outside the receptive fields of the recorded S1 neurons, neuronal sample sizes in the modulation index comparison panel were *n* = 181/199/77 cells in AIP/SMG/S1. *P*_AIP,S1_ = 0.0064, *P*_SMG,S1_ = 3.4 × 10^−^⁴; two-tailed rank-sum tests.

Extended Data Figure 7a. In the ramping task, sample sizes with valid accelerometer recording were *n* = 8/11/10 sessions for Participants 2/3/4 at each amplitude. Combined-task sessions were also included. For the measured force, Participant 2: *P*_Supra1,Supra5_ = 3.11 × 10^−^⁴ after correction; *P*_Supra2,Supra5_ = 0.015 after correction. Participant 3: P_Supra1,Supra5_ = 0.005 after correction; P_Supra2,Supra5_ = 0.018 after correction. Participant 4: P_Supra1,Supra5_ = 6.6 × 10^−^⁴ after correction; P_Supra2,Supra5_ = 0.0046 after correction. Two-tailed rank-sum tests on session means, with Benjamini–Hochberg FDR correction across the two comparisons within each participant.

Extended Data Figure 7b. Sample sizes are the same as in panel a. No sessions contained felt trials at Sub1 and catch trials in the ramping task. For the felt intensity, Participant 2: *P*_Supra1,Supra5_ = 1.6 × 10^−^⁴, P_Supra2,Supra5_ = 1.6 × 10^−^⁴ after correction. Participant 3: *P*_Supra1,Supra5_ = 3.2 × 10^−^⁴ after correction; P_Supra2,Supra5_ = 0.0064 after correction. Participant 4: *P*_Supra1,Supra5_ = 8.8 × 10^−^⁴ after correction; P_Supra2,Supra5_ = 0.009 after correction. Two-tailed rank-sum tests on session means, with Benjamini–Hochberg FDR correction across the two comparisons within each participant.

Extended Data Figure 7c. In the ramping task, M1 area comprised *n* = 186 cells, including 32 from Participant 3 and 154 from Participant 4.

Extended Data Figure 7d. For felt versus non-felt choice probability at near-threshold ramping amplitudes, Participant 3: *P*_AIP,S1_ = 0.0067 before and 0.011 after correction; *P*_SMG,S1_ = 0.0072 before and 0.011 after correction; *P*_M1,S1_ = 0.12 before and after correction. For Participant 3, fewer trials were available at near-threshold ramping amplitudes; comparisons used unpaired permutation tests with 20,000 permutations. Participant 4: *P*_AIP,S1_ = 0.0028 before and 0.0084 after correction; *P*_SMG,S1_ = 0.017 before and 0.025 after correction; *P*_M1,S1_ = 0.13 before and after correction; two-tailed rank-sum tests. Benjamini–Hochberg FDR correction was applied across three comparisons per participant.

Extended Data Figure 7e. For force–firing-rate fits on non-felt trials in Participant 2, the R² and absolute-slope panels used the same *n* = 17/173 cells in SMG/S1, each fitted using at least five non-felt trials. For R²: *P*_SMG,shuffle_ = 0.52, *P*_S1,shuffle_ = 5.0 × 10^−^⁵, *P*_SMG,S1_ = 0.0091. For absolute slope: *P*_SMG,shuffle_ = 0.80, *P*_S1,shuffle_ = 5.0 × 10^−^⁵, *P*_SMG,S1_ = 0.0035. Observed values were compared against each neuron’s shuffle mean using paired, one-tailed permutation tests; SMG versus S1 comparisons used unpaired, two-tailed permutation tests. All tests used 20,000 permutations. Counts represent positively tuned ramping spike-sorted units pooled across sessions.

Extended Data Figure 7f. For force–firing-rate fits on felt trials in Participant 4, neuronal sample sizes were n = 105/44/86/146 cells in AIP/SMG/M1/S1. The same cells were included in the R² and slope panels. For R²: P_AIP,shuffle_ = 3.6 × 10^−^⁴ before and 0.0018 after correction; *P*_SMG,shuffle_ = 0.0297 before and 0.042 after correction; *P*_M1,shuffle_ = 0.0145 before and 0.024 after correction; *P*_S1,shuffle_ = 1.97 × 10^−^¹³ before and 1.97 × 10^−^¹² after correction. Between areas: *P*_AIP,S1_ = 0.0053 before and 0.011 after correction; *P*_SMG,S1_ = 7.4 × 10^−^⁴ before and 0.0025 after correction; *P*_M1,S1_ = 0.0018 before and 0.0043 after correction. For slope: *P*_AIP,shuffle_ = 3.0 × 10^−^¹⁹ before and 1.5 × 10⁻¹⁸ after correction; *P*_SMG,shuffle_ = 3.9 × 10^−^⁹ before and 9.9 × 10⁻⁹ after correction; *P*_M1,shuffle_ = 4.5 × 10^−^¹⁶ before and 1.5 × 10^−^¹⁵ after correction; *P*_S1,shuffle_ = 6.4 × 10^−^²⁵ before and 6.4 × 10^−^²⁴ after correction. Between areas: P_AIP,S1_ = 1.2 × 10^−^⁴ before and 2.4 × 10^−^⁴ after correction; P_SMG,S1_ = 1.9 × 10^−^⁴ before and 3.1 × 10^−^⁴ after correction; P_M1,S1_ = 0.0037 before and 0.0052 after correction. Observed values were compared against each neuron’s shuffle mean using paired, one-tailed signed-rank tests; between-area comparisons used two-tailed rank-sum tests. Benjamini–Hochberg FDR correction was applied separately across ten comparisons per metric.

Extended Data Figure 7g. For high versus low felt-intensity decoding at fixed nominal amplitude in Participant 4, neuronal sample sizes were n = 555/338/348/296 cells in S1/AIP/SMG/M1. Each dot denotes one of 50 cross validation iterations, with accuracy averaged across common amplitude levels within each session and then across three contributing sessions. *P*_AIP,S1_ = 6.4 × 10^−^¹⁰ before and 9.6 × 10^−^¹⁰ after correction; *P*_SMG,S1_ = 1.1 × 10⁻¹³ before and 3.4 × 10⁻¹³ after correction; *P*_M1,S1_ = 5.0 × 10⁻⁸ before and after correction; two-tailed rank-sum tests with Benjamini–Hochberg FDR correction across three comparisons against S1.

Extended Data Figure 8a. For felt versus non-felt response latencies in Participant 2 during the near-threshold task, neuronal sample sizes were *n* = 31/179 cells in SMG/S1, with median latencies of 219/84 ms, respectively. *P*_SMG,S1_ = 2.2 × 10^−^⁴ before and after correction. For Participant 4, neuronal sample sizes were *n* = 173/201/317/308/11 cells in S1/AIP/SMG/M1/dlPFC, with median latencies of 204/211/305/224/419 ms, respectively. *P*_SMG,S1_ = 9.1 × 10^−^⁵ before and 9.1× 10^−^⁴ after correction; *P*_AIP,SMG_ = 6.4 × 10^−^⁴ before and 0.0031 after correction; *P*_SMG,M1_ = 0.001 before and 0.0035 after correction. For Participant 3, whose stimulation site was outside the receptive field of the S1 array, neuronal sample sizes were *n* = 108/283/268/159/36 cells in S1/AIP/SMG/M1/PMv, with median latencies of 214/163/192.5/210/213.5 ms, respectively. *P*_AIP,S1_ = 0.0012 before and 0.006 after correction; *P*_AIP,SMG_ = 0.0012 before and 0.006 after correction; *P*_AIP,M1_ = 0.010 before and 0.034 after correction. Two-tailed rank-sum tests with Benjamini– Hochberg FDR correction across one comparison for Participant 2 and ten pairwise comparisons within each of Participants 3 and 4. All remaining comparisons were nonsignificant after correction.

Extended Data Figure 8b. For ramping-task response latencies in Participant 3, assessed by comparing +100% stimulus and catch trials, neuronal sample sizes were *n* = 72/258/237/142/15 cells in S1/AIP/SMG/M1/PMv, with median latencies of 175/250/300/350/400 ms, respectively. *P*_S1,AIP_ = 0.017 before and 0.034 after correction; *P*_S1,SMG_ = 8.4 × 10^−^⁴ before and 0.0045 after correction; *P*_S1,M1_ = 9.0 × 10^−^⁴ before and 0.0045 after correction; *P*_AIP,SMG_ = 0.0069 before and 0.023 after correction; P_AIP,M1_ = 0.012 before and 0.031 after correction. For ramping-task response latencies in Participant 4, neuronal sample sizes were *n* = 344/269/174/224/38 cells in S1/AIP/SMG/M1/dlPFC, with median latencies of 200/200/450/250/650 ms, respectively. *P*_S1,AIP_ = 0.010 before and 0.011 after correction; *P*_S1,SMG_ = 3.1 × 10^−^²⁶ before and 3.1 × 10^−^²⁵ after correction; *P*_S1,M1_ = 4.1 × 10^−^⁷ before and 6.8 × 10^−^⁷ after correction; *P*_S1,dlPFC_ = 6.5 × 10^−^⁹ before and 1.6 × 10^−^⁸ after correction; *P*_AIP,SMG_ = 2.0 × 10^−^²³ before and 1.0 × 10^−^ ²² after correction; *P*_AIP,M1_ = 0.0077 before and 0.0096 after correction; *P*_AIP,dlPFC_ = 6.0 × 10⁻⁸ before and 1.2 × 10^−^⁷ after correction; *P*_SMG,M1_ = 5.1 × 10^−^¹⁶ before and 1.7 × 10⁻¹⁵ after correction; *P*_M1,dlPFC_ = 2.6 × 10^−^⁶ before and 3.7 × 10^−^⁶ after correction. Two-tailed rank-sum tests with Benjamini–Hochberg FDR correction across ten pairwise area comparisons within each participant. All remaining comparisons were nonsignificant after correction. Results for Participant 2 are shown in Figure 3l and are therefore omitted here. Extended Data Figure 8c–f. For felt-trial Procrustes alignment in Participant 3, neuronal sample sizes were n = 12/207/91/19/3 cells in S1/AIP/SMG/M1/PMv, respectively. For Participant 4, sample sizes were n = 22/48/202/84/11 cells in S1/AIP/SMG/M1/dlPFC, respectively. Trajectories from perception-sensitive neurons were represented by the first three principal components and aligned to S1, AIP or SMG. All 12 reference–target comparisons within each participant returned *P* = 9.99 × 10^−^⁴; one-tailed permutation tests for smaller Procrustes disparity, using 1,000 time shuffles. No shuffled disparity was as small as the observed disparity.

Extended Data Figure 8g–i. For Participant 2, neuronal sample sizes were *n* = 24/161 cells in SMG/S1, respectively. For Participant 3, sample sizes were *n* = 145/326/359/209/42 cells in S1/AIP/SMG/M1/PMv. For Participant 4, sample sizes were *n* = 22/48/202/84/11 cells in S1/AIP/SMG/M1/dlPFC.

Extended Data Figure 9a. For the behavioural psychometric fit in the discrimination task in Participant 4, *n* = 352 trials from one example session, with 32 trials at each of 11 comparison levels. The nominal standard force was 0.153 N. The logistic fit yielded a 50% response point of 0.165 N, JND = 0.0369 N, R² = 0.967.

Extended Data Figure 9b. For an example S1 neuron in Participant 4, psychometric JND = 0.0369 N; neurometric JND = 0.044 N, R² = 0.932 for neurometric fit, *n* = 352 trials. Extended Data Figure 9c. For the neurometric-to-behavioural JND comparison in Participant 4, sample sizes were n = 24/9/11/16 sensory-responsive neurons with valid fits in S1/AIP/SMG/M1. Two-tailed Welch’s t-tests versus S1, *P* before/after Benjamini– Hochberg FDR correction across three comparisons: AIP, 0.025/0.048; SMG, 0.032/0.048; M1, 0.079/0.079.

Extended Data Figure 9d. For high-versus low-force decoding at matched behavioural choice in Participant 4, pseudo-populations contained 25 neurons per area from three sessions. Trials within the psychometric JND range were resampled to match choices between force groups; each dot denotes one of 50 iterations. One-tailed Wilcoxon signed-rank tests versus chance, *P* before/after FDR correction: S1, 5.0 × 10^−^¹⁰/1.2 × 10^−^⁹; M1, 4.1 × 10^−^¹⁰/1.2 × 10^−^⁹; AIP, 0.0046/0.0062; SMG, 0.0050/0.0062; dlPFC, 0.74/0.74. Two-tailed Welch’s t-tests versus S1 across iterations, *P* before/after Benjamini–Hochberg FDR correction: AIP, 1.1 × 10^−^⁷/1.4 × 10^−^⁷; SMG, 3.6 × 10^−^¹³/1.5 × 10^−^¹²; dlPFC, 8.7 × 10^−^⁹/1.7 × 10^−^⁸; M1, 0.16/0.16.

Extended Data Figure 9e. For stronger-versus-weaker choice decoding at matched force in Participant 4, choices were balanced within each force level. Pseudo-populations contained 64 neurons per area from three sessions; each dot denotes one of 50 resampling iterations. All areas decoded above chance, one-tailed Wilcoxon signed-rank tests, *P* ≤ 7.3 × 10^−^¹⁰ before and after FDR correction. Two-tailed Welch’s t-tests versus S1 across iterations, *P* before/after FDR correction across three comparisons: AIP, 0.0055/0.0055; SMG, 0.0040/0.0055; M1, 1.8 × 10^−^⁶/5.4 × 10^−^⁶.

Extended Data Figure 9g. Example neuronal responses during report and no-report blocks. For Participant 3, report blocks included 71 felt, 34 non-felt and 45 catch trials; no-report blocks included 105 stimulus and 45 catch trials. For Participant 4, report blocks included 105 felt, 7 non-felt and 48 catch trials; no-report blocks included 112 stimulus and 48 catch trials.

Extended Data Figure 9h. For cross-session matching, Participant 3 contributed *n* = 5,703 neurons across 15 sessions, yielding 105 session pairs; mean ± s.d. match rate, 6.03 ± 3.53%; median, 5.08%; range, 1.42–17.66%. Participant 4 contributed 8,752 neurons across 16 sessions, yielding 120 session pairs; mean ± s.d., 6.94 ± 3.29%; median, 6.29%; range, 2.56–24.96%. s.d. describes variability across session pairs.

Extended Data Figure 9i. For putatively matching examples, the neuron is from AIP in Participant 3 and the two sessions are 41 days apart: waveform Pearson *r* = 0.998, KS_ISI_ = 0.017, *P* = 1 × 10^−^⁶. For Participant 4, the neuron is from M1 and the two sessions are 14 days apart: *r* = 0.998, KS_ISI_ = 0.02, *P* = 2 × 10^−^⁷.

Extended Data Figure 9j. For non-matching examples, the neurons are from M1 in Participant 3 and the two sessions are 41 days apart: waveform Pearson *r* = 0.598, KS_ISI_ = 0.738, *P* = 0.996. For Participant 4, the neurons are from AIP and the two sessions are 14 days apart: *r* = 0.319, KS_ISI_ = 0.809, *P* = 0.999. For panels i,j, *P* denotes the joint matching *P* value obtained by combining the waveform- and ISI-based empirical *P* values using Fisher’s method.

Extended Data Figure 10a. For single-area activation and multi-area coactivation during felt trials in the near-threshold task, sample sizes were n = 715 trials across 15 sessions for Participant 3 and 885 trials for Participant 4. Neuronal sample sizes were *n* = 2,247/1,328/1,831/1,710/874 cells in S1/AIP/SMG/M1/PMv for Participant 3 and 3,587/1,938/1,815/1,719/1,132 cells in S1/AIP/SMG/M1/dlPFC for Participant 4. Combined-task sessions were also included. An area was active when at least one neuron met the activation criterion. Bars show the percentage of felt trials with activation in all indicated areas. Only significant categories are shown: 12/31 for Participant 3 and 5/31 for Participant 4. Significance was assessed using one-tailed tests with 10,000 within-session shuffles of felt, non-felt and catch labels, followed by Benjamini–Hochberg FDR correction across 31 categories per participant.

Extended Data Figure 10b. For coactivation relative to independence, sample sizes were n = 688 felt trials for Participant 3 and 885 trials for Participant 4. Neuronal sample sizes were n = 2,127/1,256/1,722/1,571/806 cells in S1/AIP/SMG/M1/PMv for Participant 3 and 3,587/1,938/1,815 cells in S1/AIP/SMG for Participant 4. One-tailed permutation tests using 10,000 independent within-session shuffles of area activation: Participant 3, *P* = 0.0255; Participant 4, *P* = 0.0248. For Participant 4, the AIP + SMG + S1 + M1 combination is not shown because the observed coactivation rate was not significantly higher than expected if the four areas activated independently: observed, 62.71%; expected, 62.24%; *P* = 0.251.

Extended Data Figure 10c. For simultaneous activation across five cortical areas in Participant 3, rasters show 115 felt trials across sessions, comprising 81/79/64/64/35 distinct neurons in AIP/SMG/S1/M1/PMv. For simultaneous activation across four cortical areas in Participant 4, rasters show 346 felt trials pooled across sessions, comprising 157/180/147/185 distinct neurons in AIP/SMG/S1/M1. Each panel includes multiple trials from the same session and trials pooled across sessions. Activity on the same row across area panels was recorded simultaneously during the same trial.

Extended Data Figure 10d. For choice probability as a function of the number of cortical areas, neuronal sample sizes were *n* = 161/360/388/230/47 cells in S1/AIP/SMG/M1/PMv for Participant 3 and 235/272/475/443/24 cells in S1/AIP/SMG/M1/dlPFC for Participant 4. Mean choice probability exceeded shuffled estimates for 30 of 31 area combinations in Participant 3 (except PMv alone; FDR-adjusted *P* ≤ 0.0031) and all 31 combinations in Participant 4 (all raw and FDR-adjusted *P* = 9.99 × 10^−^⁴), based on 1,000 shuffles, with Benjamini–Hochberg correction across comparisons within each participant. Mean choice probability generally increased with the number of cortical areas, with mean slopes of 0.0205 and 0.0409 per added area in Participants 3 and 4, respectively. Both observed slopes exceeded slopes obtained from label-shuffled data; one-sided permutation tests, both P = 9.99 × 10^−^⁴.

